# The regulation of virulence gene expression is controlled by phase separation of heterochromatin protein 1 (HP1) in *Plasmodium falciparum*

**DOI:** 10.1101/2025.09.23.678141

**Authors:** D.V. Mamatharani, Devatrisha Purkayastha, Igor Niederwieser, Sandeep K. Rai, Prakshi Gaur, Mahipal Ganji, Samrat Mukhopadhyay, Till S. Voss, Krishanpal Karmodiya

## Abstract

Malaria caused by *Plasmodium falciparum* involves antigenic variation on the infected red blood cell surface by a mutually exclusive expression of virulence (*var)* genes. The repressed *var* genes have a distinct genome organization where they localize as a cluster near the nuclear periphery and are bound by PfHP1 (*P. falciparum* Heterochromatin Protein 1). However, the mode of regulation of *var* genes by PfHP1 remains unclear. In this study, we show that PfHP1 undergoes phase separation *in vitro* in an RNA and DNA-dependent manner. Single-molecule DNA tethering experiments further revealed that AT-rich DNA sequences act as nucleation sites for the assembly and compaction of PfHP1-mediated heterochromatinization. We have also identified point mutations in the IDRs (intrinsically disordered regions) of PfHP1 that disrupt its phase separation as well as DNA compaction *in vitro*. To assess the dynamic properties of PfHP1 condensates *in vivo*, we performed fluorescence recovery after photobleaching (FRAP) in GFP tagged PfHP1 parasites, which revealed rapid fluorescence recovery, supporting their fluidity and phase separation behaviour. Ectopic expression of PfHP1 phase separation and DNA compaction mutants led to dispersed nuclear localization of PfHP1, in contrast to the punctate appearance of the wild-type protein and altered chromatin binding at *var* genes. These results were corroborated with DiCre/loxP-based conditional expression of the same PfHP1 mutants, which also led to the de-repression of multiple *var* genes (as many as 54 out of 60 *var* genes), mimicking the phenotype of PfHP1 depletion mutants. Hi-C sequencing of PfHP1 mutants revealed loss of interactions in the heterochromatic regions, indicating that PfHP1 phase separation is an essential mechanism for repressive cluster formation. In conclusion, our study demonstrates the role of PfHP1-mediated phase separation in heterochromatin formation and *var* gene silencing; unveiling a fundamental mechanism that drives antigenic variation in *P. falciparum*.

## Introduction

Malaria is a protozoan disease predominantly prevalent in tropical regions. Among the five *Plasmodium* species that infect humans, *Plasmodium falciparum* is the most lethal (1). The parasite replicates within red blood cells during a phase known as the intraerythrocytic development cycle (IDC). During this cycle, a family of virulence genes called *var* genes, which encode the *P. falciparum* Erythrocyte Membrane Protein 1 (PfEMP1), are expressed. PfEMP1 proteins are exported to the surface of infected red blood cells, where they mediate cytoadherence by interacting with endothelial receptors, a key contributor to disease pathogenesis (2–4). However, the exposure of PfEMP1 on the red blood cell surface makes the parasite vulnerable to recognition by the host immune system. To counter this, the parasite has evolved a large and diverse repertoire of *var* genes, generated through recombination (5,6). Importantly, only one *var* gene is expressed at a time, a phenomenon known as mutually exclusive expression, which plays a crucial role in immune evasion and longevity of infection (7–9). The parasites also have the ability to switch the *var* gene expression and this switching delays immune recognition, providing a survival advantage (7).

The mutually exclusive expression of *var* genes is tightly regulated at the transcriptional level through epigenetic mechanisms (10). The *var* loci are highly dynamic, both temporally and spatially regulated. Their regulation is underpinned by reversible histone modifications: active marks such as H3K9ac and H3K4me3, and repressive marks like H3K9me3 (11–15). Notably, H3K9me3 and its associated heterochromatin protein, *P. falciparum* Heterochromatin Protein 1 (PfHP1), are primarily enriched at *var* gene loci and a few other clonally variant gene (CVG) families (16–18). The *var* genes, that are mostly located on the sub-telomeric regions of the chromosome are kept repressed by PfHP1 (18). They are typically cluster along with telomeres (also called as peri nuclear repressive centers, PERC) at the nuclear periphery, and upon activation, a single *var* gene repositions out of the cluster to a distinct perinuclear expression site (PES) (13,19).

HP1 is a conserved protein found across diverse eukaryotic lineages, including *Plasmodium spp*. The classical model of heterochromatin formation suggests that proteins like HP1 promote chromatin compaction and looping, thereby excluding transcriptional activators and RNA Polymerase II (20). However, the dynamic and localized nature of heterochromatin is more recently explained by the phenomenon of phase separation of HP1, observed in humans, mouse, fruit flies, and yeast (20–24). Phase separation of protein and nucleic acids allows formation of distinct, membrane-less compartments that are highly dynamic and spatiotemporally organized (25–32).

While PfHP1 is known to repress *var* gene expression (18), the molecular mechanism underlying this silencing remains poorly understood. In this study, we investigate whether phase separation of PfHP1 contributes to heterochromatin formation and gene silencing in *P. falciparum*. We employ a multi-scale approach, combining single-molecule DNA analysis, chromatin states, 3D genome organization, and reverse genetics to capture the complex regulatory landscape of *var* gene regulation. In this study, we establish a molecular model for mutually exclusive expression of *var* genes that integrates existing knowledge with our new findings.

## Results

### The *var* introns exhibit dynamic expansion and retraction of heterochromatin boundaries

It is well established that *var* gene expression occurs during the ring stage of the intraerythrocytic cycle, after which the genes enter a transcriptionally poised state (7). All *var* genes share a common structure, consisting of a large exon 1, a ∼1 kb intron, and a shorter exon 2. Notably, the intron contains an AT-rich central region and exhibits bidirectional promoter activity, giving rise to both sense and antisense transcripts (33,34).The active *var* genes, have shown to harbor H3K9ac, H3K4me3, PfH2A.Z/PfH2B.Z double histone variant (35–37). Histone modifiers such as PfSir2A, PfSir2B, Hda2 and PfSET2 were also shown to be involved in *var* regulation (13,38–41). Another mechanism involving lncRNA coming from the bidirectional promoter from the *var* intron is also in the play (34,42). We sought to investigate how these different transcriptional outputs associate with specific chromatin states.

To understand the epigenetic landscape of *var* genes, we performed chromatin immunoprecipitation followed by sequencing (ChIP-seq) for H3K9ac, H3K9me3, PfHP1, and the elongating form of RNA polymerase II (Ser2-phosphorylated) on the early ring stage parasites. Our aim was to examine the interplay of histone modifications in *var* gene regulation.

In early ring stages, we identified three distinct chromatin states at *var* loci: i) Active state - characterized by full-length sense transcripts, H3K9ac enrichment at the promoter, and widespread occupancy of RNA Pol II across the gene body (Fig. 1A). ii) Intermediate state - defined by active intronic transcription, H3K9ac enrichment at intron regions (particularly on the exon–intron junctions), and RNA Pol II peaks diverging from the intron; an indicative of bidirectional transcription (Fig. 1B). iii) Inactive state - lacking H3K9ac at both promoter and intron regions, with no RNA Pol II enrichment (Fig. 1C).

**Figure 1:**
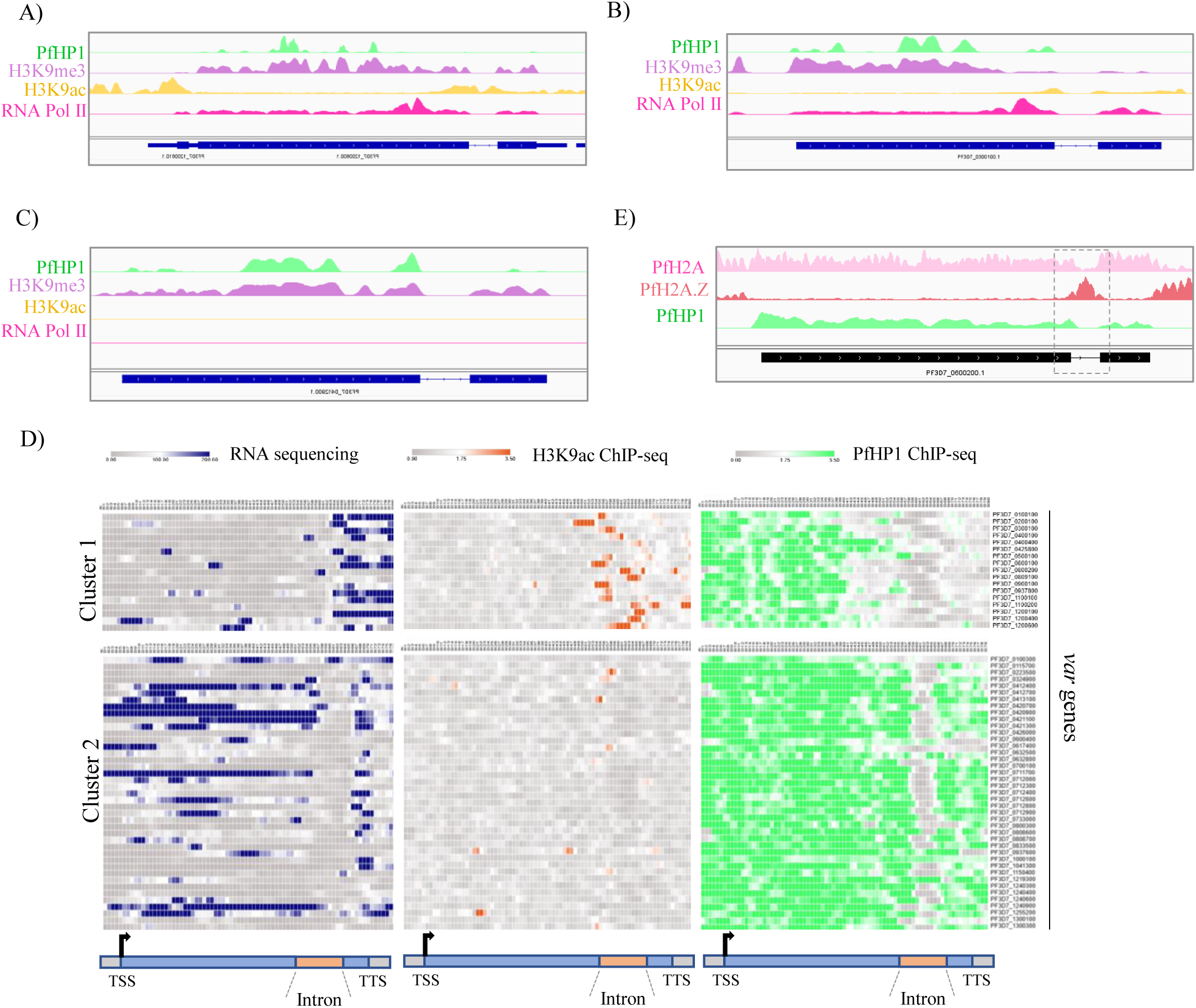
An epigenetic landscape of *var* genes. The *var* genes with ChIP sequencing peaks (input normalized) for enrichment of H3K9ac (yellow), RNA Pol II (pink), heterochromatic mark H3K9me3 (lilac) and PfHP1 (green) in early ring stage, A) active gene, B) intermediate state, and C) inactive gene. D) Mean tag density of RNA sequencing reads, H3K9ac reads (input normalized), and PfHP1 reads (input normalized) on *var* gene body. The clustering is based on presence and absence of H3K9ac on introns. The bottom schematic represents *var* gene body with two exons (blue) and an intron (peach). The color bar at the top represents the relative mean tag density for the RNA seq and ChIP seq. E) A representative example of *var* gene showing, ChIP sequencing peaks (input normalized) for enrichment of PfH2A, PfH2A.Z and PfHP1. The box highlights the intronic part. The PfH2A and PfH2A.Z ChIP-seq data is re-anlaysed from Bartfai et. al., 2010 (35).

To further see how transcriptional output from the *var* introns correlated to the histone modification and modifiers, we further performed a ChIP-seq along RNA-seq (with corresponding RNA samples) in late ring stage parasites. The analysis on this data revealed that the majority of *var* genes are devoid of heterochromatin in the intron (cluster-2, Fig. 1D). However, a subset (cluster-1) showed intronic H3K9ac enrichment and expression of sense transcripts originating from the intron. These cluster-1 genes displayed a retracted heterochromatin boundary at *var* introns compared to cluster-2. Interestingly, *var* introns contain a previously reported sequence motif with potential insulator activity (43). In cluster 1 *var* genes, activating histone marks - H3K9ac, H4K8ac, and H3K79me2 - were enriched around this motif (Supplementary Fig. 1A). In contrast, when H3K9ac was absent at this intronic motif, heterochromatin extended into this region (Supplementary Fig. 1B). The heterochromatin markers PfSir2A and H3K36me3 showed similar patterns of expansion and retraction, mirroring PfHP1.

The apparent exclusion of heterochromatin from the majority of *var* introns suggests the presence of an active mechanism that restricts its spread. One hypothesis is that these regions contain nucleosomes with different histone variants, which may limit heterochromatin propagation (44). Supporting this, we found that PfH2A.Z (data re-analyzed from Bartfai et. al., 2010), a histone variant known to mark regulatory elements (35,45), is consistently enriched at *var* introns. Notably, PfHP1 and PfH2A.Z exhibit opposing patterns of enrichment, both globally and locally at *var* loci (Fig. 1E and Supplementary Fig. 1C). This observation is consistent with findings in *Saccharomyces cerevisiae*, where H2A.Z acts as an insulator to the spread of heterochromatin (46). Thus, PfH2A.Z, possibly in concert with other insulator factors, may contribute to the establishment of intronic chromatin boundaries.

To further explore chromatin structure at *var* introns, we reanalyzed ATAC-seq data from Toenhake et al., 2018 (47) and observed increased chromatin accessibility in these regions. Coupling this with strand-specific RNA-seq data from the same study revealed that many *var* introns produce divergent transcripts (Supplementary Fig. 1D), suggesting a correlation between open chromatin and transcriptional activity at the intron. Although heterochromatin generally envelops *var* loci, intronic regions exhibit dynamic boundary shifts. The complex interplay between epigenetic modifications, chromatin-associated proteins like PfHP1, and transcriptional dynamics presents challenges in fully resolving the mechanism of heterochromatin mediated regulation. We hypothesize that phase separation of PfHP1 is one such mechanism that might integrate highly dynamic spatiotemporal regulation of *var* gene expression.

### PfHP1 undergoes phase separation *in vitro*

Given the central role of PfHP1 in defining chromatin states at *var* loci, we next sought to investigate its intrinsic biophysical properties that might contribute to heterochromatin organization. In particular, we examined whether PfHP1, like its homologs in other systems, is capable of undergoing phase separation, a mechanism increasingly recognized for organizing chromatin domains in a dynamic and reversible manner. Although HP1 homologs across species are known to undergo phase separation, sequence comparisons revealed that PfHP1 shares limited similarity with its counterparts (Supplementary Fig. 2A). Nonetheless, PfHP1 retains the canonical chromodomain (CD) and chromo shadow domain (CSD), connected by a highly disordered hinge region (16). Both domains contain conserved amino acid stretches across species (Supplementary Fig. 2B), suggesting that despite sequence divergence, HP1 proteins may employ a shared biophysical mechanism, such as phase separation, in chromatin regulation.

To further investigate this, we performed disorder prediction analysis, which revealed that multiple regions of PfHP1, including the N-terminal extension (NTE), hinge, CSD, and C-terminal extension (CTE) harbor intrinsically disordered regions (IDRs) (Supplementary Fig. 2C), a known feature of phase-separating proteins (25). We then examined the droplet formation potential of PfHP1 *in vitro*. Both untagged and GFP-tagged PfHP1 readily formed droplets (Fig. 2A and Supplementary Fig. 2D,E). Droplet formation was concentration-dependent, with saturation concentrations (C_sat_) of 37.4 μM for untagged PfHP1 and 11.54 μM for PfHP1-eGFP, suggesting that GFP tagging modestly enhances phase separation (Fig. 2B and Supplementary Fig. 2F). The presence of a molecular crowding agent (PEG8000) reduced the C_sat_ to 28.5 μM for untagged PfHP1 (Supplementary Fig. 2E,F). The PfHP1 condensates displayed hallmark features of liquid-like behavior, such as droplet fusion, surface wetting, and dripping (Supplementary Video 1). Moreover, fluorescence recovery after photobleaching (FRAP) assays showed rapid signal recovery, indicating high internal mobility within droplets (Fig. 2C and Supplementary Fig. 2G,H). Finally, to assess the forces driving this condensate formation, we tested salt sensitivity. Increasing salt concentrations beyond 200 mM disrupted droplet formation, highlighting the importance of electrostatic interactions in maintaining phase-separated states (Supplementary Fig. 2I). Together, these findings establish that PfHP1 possesses intrinsic biophysical properties required for phase separation. However, in cells the formation of such biomolecular condensates is governed by weak multivalent interactions between the proteins and nucleic acid (28,32).

**Figure 2:**
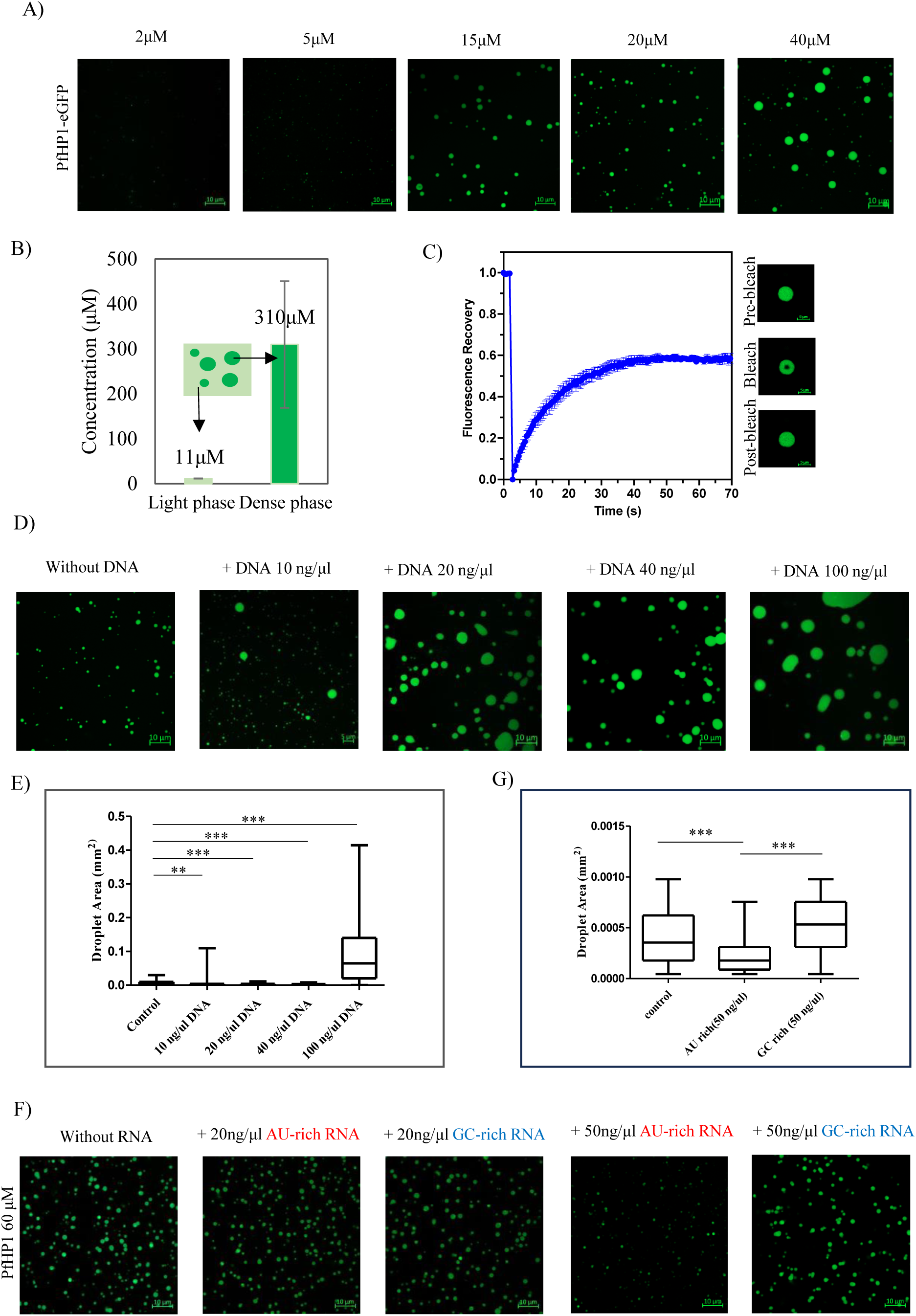
PfHP1undegoes phase separation *in vitro*. A) Concentration dependent assembly of PfHP1-eGFP condensates. The concentration of the protein used are indicated on the top of each image. The reaction buffer used was 20mM HEPES pH 7.4, 150mM KCl (scale bar, 10 μm). B) Saturating concentration (C_sat_) estimation of the PfHP1-eGFP (in reaction buffer, 20 μM protein). The concentration of the light phase is C_sat_ and the concentrations are indicated on the top of each bar graph. C) FRAP assay showing recovery of PfHP1-eGFP (means ± SD; n = 5). The microscopy images of droplet before the photo bleaching, after the bleaching and recovery are given at the bottom of the graph for reference (scale bar, 5 μm). D) Droplet assay with PfHP1-eGFP with increasing concentration of parasite DNA (in reaction buffer, 20 μM protein). The concentrations of DNA used is indicated on the top of each image (scale bar, 10 μm). E) The quantification of droplet size with different DNA concentrations (control: n=82, 10 ng/μl DNA: n=283, 20 ng/μl DNA: n=57, 40 ng/μl DNA: n=46, 100 ng/μl DNA: n=30). The statistical significance was calculated using non-parametric t-test (significance value summary is represented as asterisk on top of the graph). F) The microscopy snap shots of droplet assay with PfHP1 and IVT ncRNAs (GC-rich and AU-rich). Two different concentrations of RNA were used 20 ng/μl and 50 ng/μl along with 60 μM PfHP1, 1% S33C_PfHP1-Alexa fluor 488 (in reaction buffer, scale bar, 10 μm). G) Quantification of droplet area with AU-rich and GC-rich ncRNAs, the number of droplets used for the quantification are 251 for control, 92 for AU-rich RNA, and 209 for GC-rich RNA. The statistical significance was calculated using non-parametric t-test (significance value summary is represented as asterisk on top of the graph).

### PfHP1 phase separation exhibits tunable sensitivity to nuclear components

Given that PfHP1 can undergo phase separation *in vitro*, we next explored whether this behavior is influenced by the molecular environment of the nucleus. Nucleic acids, particularly DNA and RNA, constitute a major component of the nuclear milieu and are known to modulate phase transitions of nuclear proteins (21,48,49). HP1 homologs are reported to bind DNA and compact chromatin in a cooperative manner, and DNA has been shown to promote HP1 phase separation in other systems (21,50). While RNAs play important roles in heterochromatin nucleation and maintenance, their influence on HP1-mediated biomolecular condensate formation is not known in any system (51,52). To investigate these possibilities, we tested whether DNA and RNA could modulate PfHP1 droplet formation *in vitro*.

We observed that increasing concentrations of a *P. falciparum* genomic DNA fragment (800 bp) promoted droplet formation by PfHP1-eGFP (Fig. 2D,E and Supplementary Fig. 3A), and slightly increased the rate and extent of FRAP recovery (Supplementary Fig. 3B), indicating more dynamic internal rearrangements. Similar results were obtained using 146 bp Widom 601 DNA (identified by Lowary and Widom using the SELEX method (53)), where the increasing concertation of DNA promoted the phase separation (Supplementary Fig. 3C). Additionally, DNA lowered the critical concentration required for droplet formation (Supplementary Fig. 3D). Fluorescent labeling of 601 DNA with YOYO-1 dye confirmed its incorporation into PfHP1 droplets, suggesting that DNA acts not merely as a crowding agent but is physically integrated into the condensates (Supplementary Fig. 3E). These findings suggest that genomic DNA may facilitate the PfHP1 phase separation and that *var* genes may be compartmentalized by the phase separated droplet.

We next tested whether RNA could similarly modulate PfHP1 droplet formation. *P. falciparum* telomeric and sub-telomeric regions produce numerous AU-rich non-coding RNAs, reflecting the AT-rich genome composition (54,55). Using *in vitro* transcribed AU-rich and GC-rich RNA, we found that AU-rich RNA caused dose-dependent dissolution of PfHP1 droplets (Fig. 2F,G), while GC-rich RNA had no observable effect in the tested concentration range. These results suggest that AU-rich RNA may destabilize PfHP1 condensates, potentially serving as a regulatory mechanism. Lastly, we examined the influence of PfH2A.Z on PfHP1 condensate formation. While control histone PfH2A caused only a modest reduction in droplet formation, addition of PfH2A.Z led to complete dissolution of PfHP1 droplets (Supplementary Fig. 3F), consistent with the antagonistic relationship between PfHP1 and PfH2A.Z observed on chromatin. Collectively, these findings reveal that nucleic acids and chromatin-associated factors can dynamically modulate PfHP1 phase separation, potentially contributing to the regulation of heterochromatin domains in *P. falciparum*. The IDRs are known to engage in weak multivalent interactions with nucleic acids and other proteins; hence, we next tried to understand how these aspects contribute to the phase separation of PfHP1.

### Point mutations in the intrinsically disordered regions of PfHP1 disrupts phase separation

Next, we tried to identified the point mutations in PfHP1 that disrupts its multivalent interactions and oligomeric property. Based on the limited literature in *P. falciparum* it was hard to probe for such biochemical properties. Hence, mutations were chosen based on conserved residues in the chromodomain (CD) and chromo shadow domain (CSD) of PfHP1 and Swi6, a well-characterized HP1 homolog from *Saccharomyces pombe*. Although PfHP1 shares limited overall sequence similarity with Swi6, many residues within its CD and CSD were conserved (Supplementary Fig. 3G). Previous studies on Swi6 identified key functional mutations, such as LoopX (a mutation in K94A in the ARK loop that disrupts oligomerization), CageX (W104A), AcidX (E(74–80)A), (both of which impairs binding to H3K9me3), and DimerX (L315D, which disrupts homodimerization) and amongst these mutations LoopX and DimerX affect its phase separation property (22,56,57). We hypothesized that analogous mutations in PfHP1 would have similar effects.

Using pairwise sequence alignment, we identified corresponding residues in PfHP1 and generated four mutants via site-directed mutagenesis: E6A/E7A/E9A (AcidX equivalent, termed E-A), K17-20A (lysine stretch mutation, K-A), W29A (CageX equivalent, W-A), and L254A/D (DimerX equivalent, L-A and L-D) (Fig. 3A and Supplementary Fig. 3G). Notably, we did not detect a canonical ARK loop in PfHP1’s chromodomain, but a cluster of four lysines likely plays a role in multivalent interactions such as oligomerization or DNA binding through their positive charge. Given the role of IDRs in multivalent interactions and phase separation, we noted that, all four mutations were located within predicted IDRs of PfHP1 (Supplementary Fig. 2C and Fig. 3A). The sequence analysis of wild-type and mutant PfHP1 predicted that E-A and K-A would not phase separate, whereas W-A and L-D would (Fig. 3A). Functional assays revealed that the AcidX mutant E-A indeed failed to form droplets under conditions where wild-type PfHP1 readily phase separated (Fig. 3B). However, droplet formation was partially rescued by adding *P. falciparum* DNA or the crowding agent PEG8000, suggesting that the mutant’s critical concentration to phase separation is higher and can be overcome by lowering the threshold by addition of crowding agent such as PEG. The K-A mutant, targeting the lysine-rich region, showed the most severe phenotype, completely abolishing droplet formation with no rescue by DNA or PEG (Fig. 3B). The CageX mutant W-A also lacked phase separating property at usual concentrations but showed partial recovery with PEG8000, similar to E-A (Fig. 3B). In contrast, the two dimer mutants, L-A and L-D, surprisingly had no detectable effect on phase separation despite the L-D mutation introducing a polar charged residue instead of a hydrophobic leucine (Fig. 3C). Together, these results highlight the critical roles of specific conserved residues and regions within PfHP1 in driving phase separation, and underscore that disruption of key interactions, especially involving charged residues, can strongly impair phase separation.

**Figure 3:**
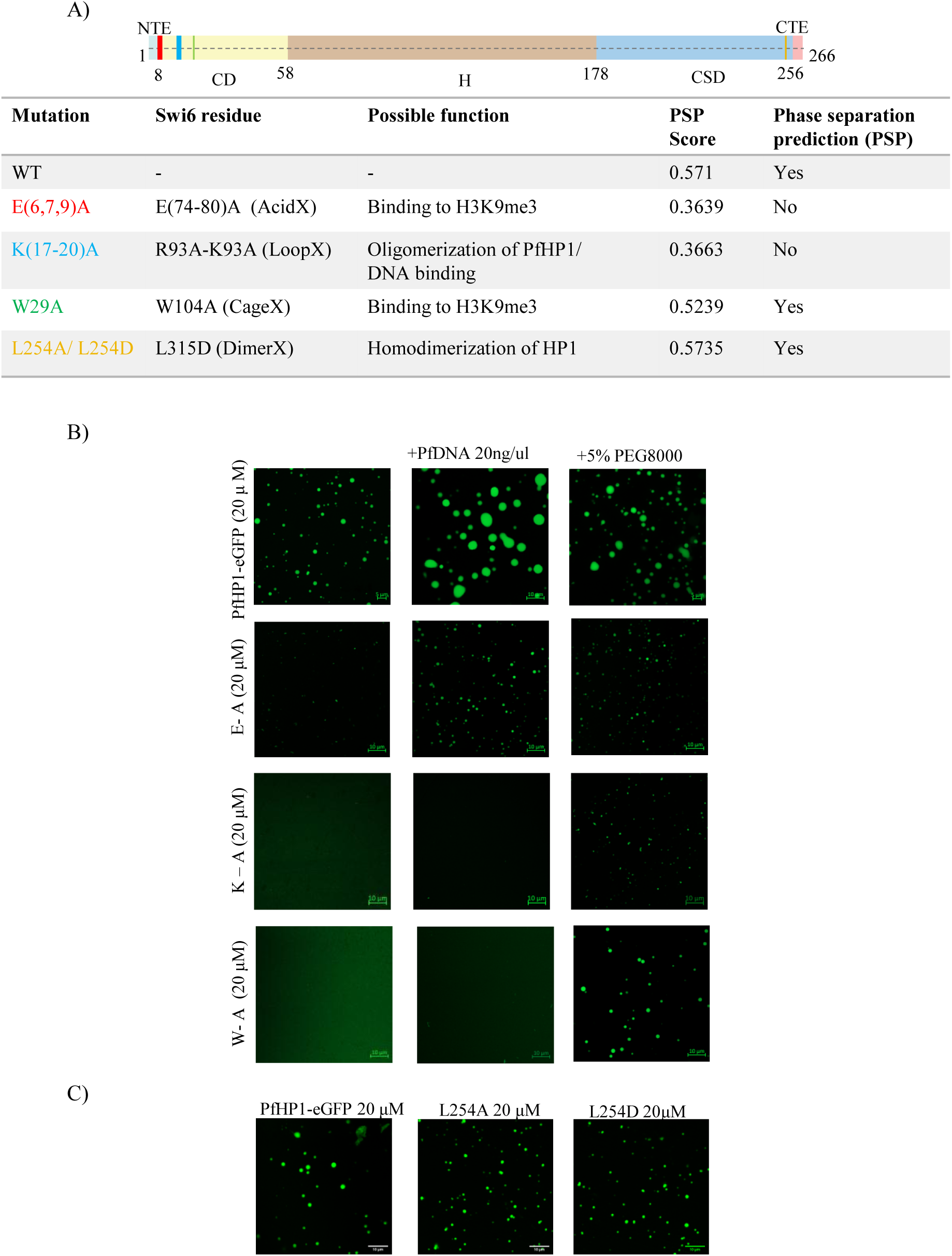
Droplet assay with PfHP1 mutants. A) A schematic representation of PfHP1 mutants (E-A: red, K-A: blue, W-A: green, L-D: yellow) and their predicted functions based on pairwise alignment to Swi6 are summarized as a table. The table also includes the phase separation prediction for each mutant using PSPred. A PSP score greater than 0.5 represensts propensity to phase separate. B) Droplet assay with PfHP1-eGFP mutants with mutations in NTE (E-A) and CD (K-A, W-A) (20 μM of each protein in the reaction buffer 20mM HEPES pH 7.4, 150mM KCl was used). Used 5% PEG8000 and 20ng/μl PfDNA in the reaction buffer to see if any recovery. C) Droplet assay with PfHP1-eGFP mutant with L to D or L to A mutation in the CSD (20 μM of each protein in the reaction buffer); PfHP1-eGFP wild-type was used as a control in all the assays.

### Single molecule imaging reveal PfHP1 forms co-condensates and compacts AT-rich *var* intronic DNA

Having established that specific conserved residues within PfHP1 are critical for its phase separation properties, we next examined how these properties relate to its chromatin-binding preferences, particularly its dynamics on atypically AT-rich genome of *Plasmodium falciparum* (∼80%). PfHP1 exhibits a distinctive distribution, localizing primarily to sub-telomeric coding regions and telomeric regions of the genome (18). On average, heterochromatic genes show ∼70% AT content, with sharp spikes reaching 90–100% at the start and end of the coding sequence (Supplementary Fig. 4A). Notably, *var* genes contain highly AT-rich introns (∼90%), where PfHP1 shows expansion/retraction of boundaries, and comparatively less AT-rich exons (∼70%) (Supplementary Fig. 4A). Consistently we observe, for *var* genes, both promoters and introns are markedly AT-rich (Supplementary Fig. 4B–E). In contrast, euchromatic genes tend to have an overall AT content of 80–90%, strikingly higher compared to the heterochromatic genes (Supplementary Fig. 4A).

We found that PfHP1 directly binds to AT-rich DNA sequences in electrophoretic mobility shift assays (Supplementary Fig. 5A). Additionally, as we observed in the above section, dynamic changes in PfHP1 enrichment at the highly AT-rich *var* intronic regions (Fig. 1D). To investigate PfHP1 dynamics on an AT-rich substrate in real time, we employed a single-molecule DNA tethering assay using fluorescently labeled PfHP1-GFP and DNA substrates (see the experimental design in the Supplementary Fig. 5B) (58). The DNA constructs with distinct AT-richness were immobilized on a PEG-passivated slide: a 19 kbp test fragment containing a 1 kbp *var* intron with 86% AT-content in the middle of a linearized GC-rich plasmid, and an 18 kbp control fragment (just the linearized GC-rich plasmid). We double tethered the DNA by using a biotin-neutravidin interaction and visualized under a TIRF microscope. Upon introducing PfHP1-eGFP into the flow cell, we observed prominent puncta formation in the center and in some cases on an end of the 19kbp DNA, which otherwise appeared homogenous (Fig. 4A and Supplementary Video 2,3). Whereas, the control DNA exhibited only weak, unstable puncta mostly on one end of the substrate (Supplementary Fig. 5C and Supplementary Video 4,5).

**Figure 4:**
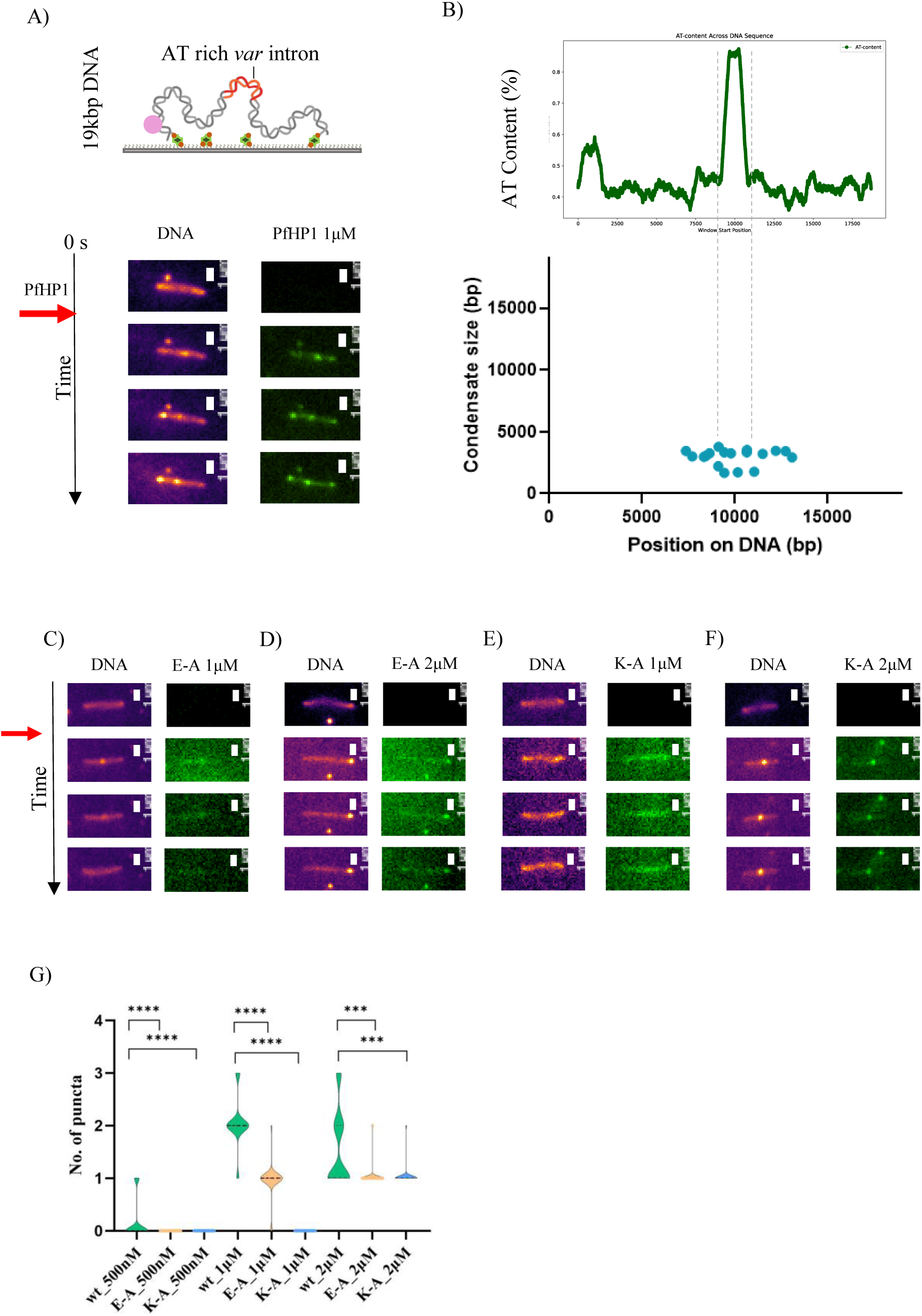
DNA tethering experiments with PfHP1-eGFP. A) Snapshots from the time series of double labelling experiment done with 19 kbp DNA with 1kbp *var* intron in the middle and PfHP1-eGFP (1 μM). The schematic at the top represents a double tethered DNA with an AT-rich intronic stretch in the middle; the DNA tethering was done by neutravidin (green) and biotin (orange) interaction; for determining directionality Cy5 (magenta) probe is attached on one end (scale bar, 1 μm; The red arrow indicates the time point at which the protein was added during the experiment). B) The quantification of the position of the puncta in the 19 kbp substrate (bottom) and AT-richness of the substrate (top) indicting a moderately AT-rich stretch at the beginning and a highly AT-rich stretch at the middle, that is *var* intron (n=15). C) Snapshots from the time series of double labelling experiment done with 19 kbp DNA and PfHP1 mutants E-A (1μM), D) E-A (2μM), E) K-A (1μM), F) E-A (2μM) (scale bar, 1 μm; The red arrow indicates the time point at which the protein was added during the experiment). G) Violin plot showing quantification of number of puncta in each condition (as indicted in the graph) used for mutants E-A and K-A. The wild-type PfHP1 was used as a control in each condition. The schematics are made in biorender.com.

Interestingly, PfHP1 initially coated the entire 19 kbp DNA strand and eventually formed stable puncta near the center, aligning with the location of the highly AT-rich intron (Supplementary Video 2,3). Using Cy5-labeled DNA, we further mapped the relative position of these puncta and found them concentrated around the 10 kbp mark, coinciding with the AT-rich sequence (Fig. 4B and Supplementary Fig. 5D). Based on the extent of compaction observed, we estimate that PfHP1 condenses approximately 2,000 bp of DNA, centered on the AT-rich region (Fig. 4B). At higher protein concentrations (1 μM and 2 μM), multiple puncta formed along individual DNA molecules (Supplementary Video 2,3 and Fig. 4G).

### Disruption of PfHP1 phase separation alters its co-condensation with AT-rich DNA

To further probe the relationship between PfHP1’s phase separation ability and its DNA binding behavior, we examined the co-condensation properties of polar residue mutants E-A and K-A using the single-molecule DNA assay. The E-A mutant, which mimics the AcidX mutation, failed to form any condensates at 500 nM. At higher concentrations, it did form co-condensates, but these were unstable even at 1 μM (Fig. 4C,G). Notably, at 2 μM, the mutant did not produce multiple puncta as observed with the wild-type protein. Instead, an unstable puncta was formed and moved along the DNA (Fig. 4D,G and Supplementary Video 6,7), suggesting a loss of sequence-dependent condensation capacity.

The K-A mutant, targeting a positively charged lysine-rich stretch, also showed no condensate formation at 500 nM and 1 μM (Fig. 4E,G). However, at 1 μM, the mutant was still able to coat the DNA (Fig. 4E and Supplementary Video 8,9). At 2 μM, K-A formed a single punctum per DNA molecule, unlike the multiple puncta typically formed by wild-type PfHP1 (Fig. 4F,G). These findings indicate that electrostatic interactions mediated by the lysine-rich region are likely critical for multivalent interactions and robust condensate formation. To quantify these effects, we calculated the proportion of double-tethered DNA molecules exhibiting puncta. Strikingly, the mutants displayed a binary behavior, either forming a punctum or failing to form any at all, highlighting the sensitivity of PfHP1 condensation to specific structural features (Supplementary Fig. 5E).

Together, these results show how specific sequence features and conserved residues within PfHP1 coordinate to drive a sequence-dependent condensation on chromatin, offering a mechanistic framework for understanding the nucleation and spread of heterochromatin in *P. falciparum.* We propose a model wherein co-condensation between PfHP1 and highly AT-rich *var* promoter or intronic regions may act as nucleation sites for heterochromatin assembly (Supplementary Fig. 5F).

### Ectopic expression of PfHP1 phase separation mutants reveals chromatin binding defects in *P. falciparum*

Following our *in vitro* characterization of PfHP1 and its phase separation-deficient mutants, we next sought to investigate their behavior in *P. falciparum* cells. Since PfHP1 is essential for parasite survival (18,59), direct mutagenesis at the endogenous locus posed a risk of lethality. Therefore, we initially adopted an ectopic expression strategy, introducing mutant constructs into a dispensable genomic locus known as *cg6* (PF3D7_0709200) via CRISPR/Cas9-based genome editing (60) (Supplementary Fig. 6A). Using this approach, we successfully generated parasite lines expressing GFP-tagged versions of wild-type PfHP1, as well as the K-A, W-A, and L-D mutants, in addition to the endogenous PfHP1. PCR on the genomic DNA, Western blot and live cell fluorescence microscopy confirmed the expression of the GFP-tagged proteins (Supplementary Fig. 6B-E). The wild-type and K-A proteins localized to discrete nuclear puncta, consistent with the expected heterochromatin foci (16,18). In contrast, the W-A and L-D mutants exhibited a more diffuse nuclear distribution and significantly weaker GFP signals, which was confirmed by reduced expression levels observed by Western blot (Supplementary Fig. 6D,E).

To assess chromatin association, we performed ChIP-seq experiments using an anti-GFP antibody. All PfHP1 mutants displayed reduced chromatin binding compared to the wild type, with W-A showing particularly low tag density (Supplementary Fig. 6F,G). Despite these defects in chromatin binding, overexpression of the mutants did not lead to noticeable growth defects or phenotypic changes in the parasites. This is likely due to the continued presence of endogenous wild-type PfHP1, which may compensate for the mutant proteins. To directly assess the functional consequences of the mutations without interference from the wild-type protein, we next aimed to introduce the mutations at the endogenous *pfhp1* locus.

### PfHP1 shows *in vivo* phase separation in *P. falciparum*

We began with a CRISPR-Cas9 strategy to replace the endogenous *pfhp1* with mutant alleles. However, repeated attempts failed to generate viable clones, suggesting that the mutations may be detrimental to the parasites. We then adopted a previously reported conditional expression system using a DiCre-loxP-based strategy (Fig. 5A) enabling temporal control of mutant expression (61–64). In this setup, endogenous PfHP1 was tagged with mScarlet-I, while the mutant version was tagged with GFP. Upon rapamycin treatment, DiCre-mediated recombination excised the wild-type allele, allowing expression of only the mutant PfHP1-GFP (Fig. 5A). We employed a one-step CRISPR/Cas9 strategy to knock-in the entire stretch at once. We used a previously described self-processing gRNA for targeting start and end of the *pfhp1* locus (65).

**Figure 5:**
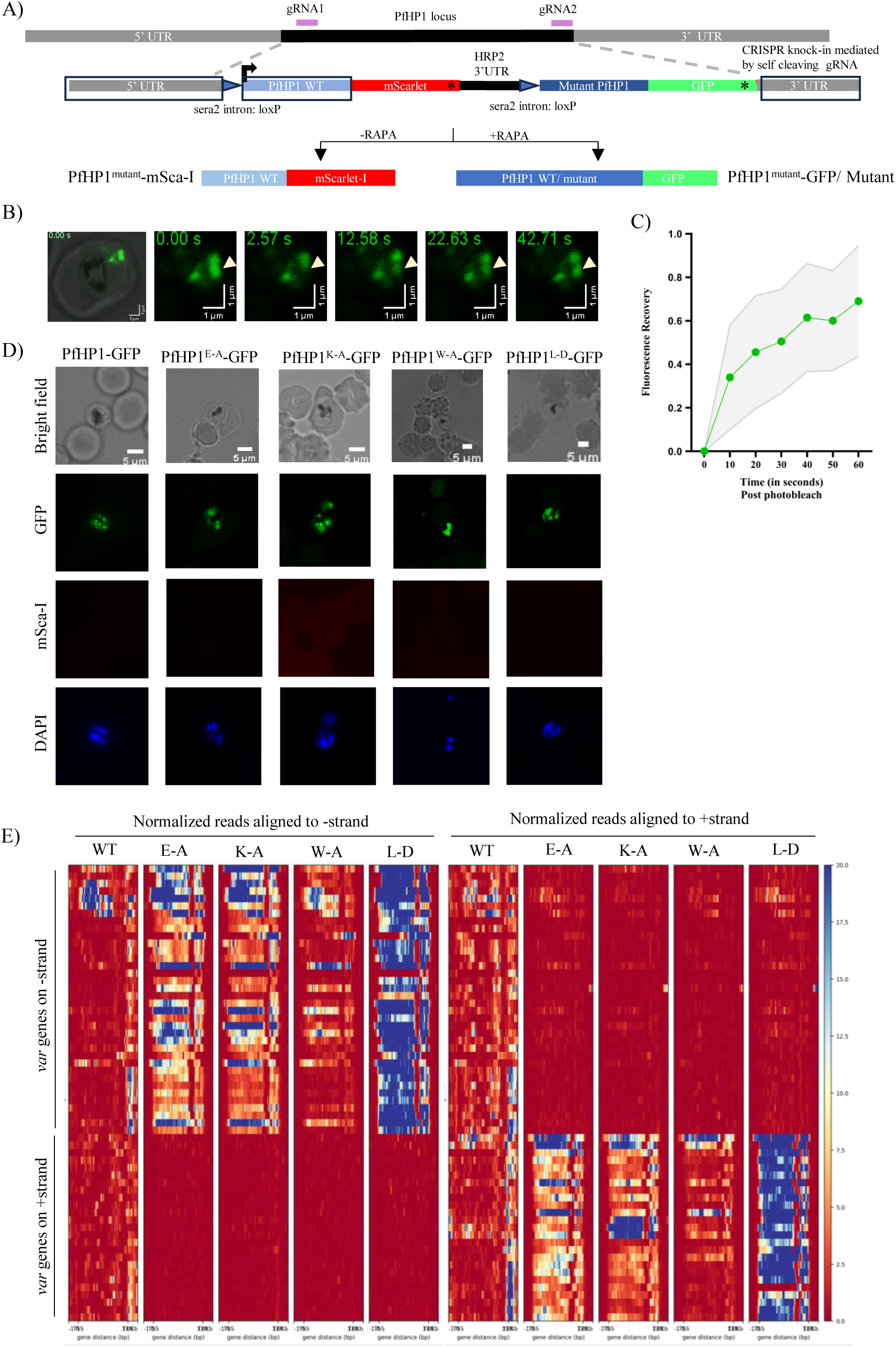
PfHP1 FRAP assay in cells and PfHP1 conditional mutant lines. A) Schematic showing one-step CRISPR-Cas9 based strategy with self-cleaving gRNAs targeting two loci for engineering DiCre-inducible PfHP1 mutant lines. Upon rapamycin treatment, the DiCre enzyme dimerizes and recombines the loxP sites to swap the endogenous wild-type *pfhp1-mscarlet-i* gene with a mutated *pfhp1-gfp* version inserted downstream. The black asterisk represents the stop codons, the purple lines are gRNAs and the three homology regions are boxed (black). B) The first image represents the DIC image of RBC with the parasite tagged for PfHP1. Representative snapshots of a FRAP assay on parasites expressing GFP-tagged wild-type PfHP1. The white arrows represent the puncta that was bleached and recovered later (scale bar, 1 μm). C) Quantification of FRAP recovery (means ± SD; n = 5). D) Live cell fluorescence imaging of mutant lines expressing PfHP1 mutants tagged with GFP upon rapamycin treatment. DAPI was used for nuclear staining (scale bar, 5 μm). E) Heat maps representing the normalized RNA reads on the gene body of all the *var* genes. The *var* genes are arranged based on their strand for ease of understanding. The sense reads for – strand *var* genes will come from the reads mapped to – strand and the same applies to the + strand. The antisense reads for the – strand genes will come from the reads mapped to the + strand. The colour bar represents relative read density. The RNA-seq experiments are done in triplicates and here we show only replicate 1 for representative purpose.

We generated a control line that expresses wild-type PfHP1 in both conditions with or without rapamycin treatment (Fig. 5A). This line expresses mScarlet-I tagged wild-type PfHP1 and upon rapamycin treatment it expresses GFP tagged wild-type PfHP1.

To probe condensate dynamics *in vivo*, we performed FRAP experiments on GFP-tagged PfHP1 puncta, which showed rapid fluorescence recovery in the timescale of seconds (Fig. 5B), indicative of liquid-like behavior. The quantification of the same showed ∼60% of recovery (Fig. 5C). This experiment was majorly challenged by the size of the PfHP1 foci that are smaller than the limit of resolution of most the microscopes (∼ 200 nm). Also, the quantification was constrained by the intrinsic property of the endoparasite, to move in both XY and Z axes, even though we immobilized the infected RBCs by poly-L-lysine on a coverslip. This results in a not very smooth recovery curve. Nevertheless, these results indicate that PfHP1 is a dynamic component of the heterochromatin, and *P. falciparum* heterochromatin has fluid nature that supports phase separation behavior rather than a static-dense structure.

### PfHP1 phase separation is essential for nuclear puncta formation and affects gene silencing

Using the DiCre/loxP approach described above, we successfully generated conditional mutant lines for E-A, K-A, W-A, and L-D, along with a wild-type PfHP1 control line. The PCR and Sanger sequencing validation of the mutant lines showed knock-in with the correct insertions (Supplementary Fig. 7A-D). The expression of mScarlet-I tagged PfHP1 confirmed the presence of the wild-type protein (Supplementary Fig. 8A), and upon rapamycin treatment, GFP-tagged mutant expression was observed (Fig. 5D). The rapamycin treatment was further validated using the Sanger sequencing to show the right recombination (Supplementary Fig. 7D).

We next examined the localization of the mutant PfHP1 proteins in live schizonts of the same generation (gen1) following the rapamycin-induced DiCre-dependent swap of the endogenous wild-type allele with the mutant allele. While wild-type and K-A mutants formed compact nuclear puncta, E-A and W-A mutants displayed diffuse localization with no puncta, similar to the results obtained with the ectopic overexpression. The L-D mutant formed puncta, though these were larger and more dispersed than wild-type PfHP1-GFP (Fig. 5D). Interestingly, the K-A mutation, which showed complete loss of phase separation *in vitro*, initially (gen1) exhibited a punctate localization in live cells. However, when analyzed in the subsequent generation (gen2), the K-A mutant displayed disrupted puncta formation (Supplementary Fig. 8B) and was lethal to the parasites (data not shown). Similarly, the E-A mutant failed to grow beyond gen2 and exhibited sexual conversion and developed into gametocytes (Supplementary Fig. 8C). The L-D mutant also displayed a lethal phenotype after two to three generations of mutant induction. The W-A mutant grew very slowly upon rapamycin treatment, suggesting growth defects potentially linked to defective merozoite formation or invasion, an aspect requiring further investigation. Some of these mutants displayed phenotypic affects as that of PfHP1 knock-down or knock-out (18,59). More specifically the ultra-high sexual conversion rates and gametocyte production in gen1 due to the de-repression of AP2-G (a key to the transcriptional switch for sexual conversion) and in gen2 asexual stages get arrested/die at trophozoites (18,59,66).

To assess the effect of PfHP1 mutants on transcription, we performed directional RNA sequencing experiment with rapamycin treatment given to the early ring (∼6-10 hpi) stage parasites and harvested after 48 hours (early rings) in triplicates in gen2 of mutant lines E-A, K-A, W-A and L-D. Rapamycin treated GFP-tagged wild-type PfHP1 line served as the control. Gene ontology analysis of upregulated genes in the mutants revealed enrichment for cytoadherence-associated genes and antigenic variation (Supplementary Fig. 9C-F). All four mutants showed global de-repression of *var* genes. Unlike the control line, which exhibited mostly intronic *var* transcripts, mutants produced full-length *var* transcripts (Supplementary Fig. 9A,B). There is a strong upregulation of multiple *var* genes in the mutant lines and a marked reduction in intronic transcription (Fig. 5E).

The degree of *var* gene activation varied across mutants: L-D showed upregulation of 54 *var* genes, E-A upregulated 24, while K-A and W-A upregulated 11 and 10 respectively (Supplementary Fig. 9G,H). These results suggest that disruption of PfHP1-driven compartmentalization leads to a transcriptionally permissive state. Whether this hyperactivation also involves repositioning of *var* loci within the nucleus remains to be explored.

In addition, other heterochromatic genes were also upregulated in the mutant lines. Notably, AP2-G, a key AP2 factor associated with the gametocytogenesis (18,66), was upregulated in all mutants (Supplementary Fig. 10A–D). Clonally variant gene families such as rifin, stevor, surfin, and phist also showed increased expression. The L-D mutant displayed the biggest impact, with the highest number of deregulated CVGs.

Together, these findings demonstrate that PfHP1 phase separation is essential for maintaining transcriptional repression across heterochromatic regions. Its disruption leads to widespread activation of *var* genes, other CVGs and AP2-G, thereby compromising the epigenetic regulation critical for parasite survival and pathogenesis. The similar type of transcriptional de-repression was observed in PfHP1 knock-down lines in earlier studies, indicating that PfHP1 phase separation mutants are disrupting the very essential function of the protein (18).

### PfHP1 phase separation helps in genome organization and heterochromatic gene clustering

Next, we tried to see if the disruption of PfHP1 phase separation leads to global changes in genome organization and *var* gene clustering in the 3D space. We employed a Hi-C sequencing approach using two restriction enzymes DpnII and HnfI that helped us attain ∼1kbp resolution (see Supplementary file 1 for the cut sites). We harvested rapamycin treated early ring stage parasites (similar experimental design as RNA seq harvesting) from phase separation mutant DiCre lines E-A, K-A and W-A as well as PfHP1-GFP, as a wild-type control.

We observed strong boundaries between the euchromatic genes and heterochromatic regions containing *var* genes (Fig. 6A). As reported earlier we observed high contact frequency at centromeres, heterochromatin and telomers across all the chromosomes (Supplementary Fig. 11) (67,68). The internal heterochromatin islands showed interaction with itself and sub-telomeric heterochromatin (Fig. 6A). As discussed earlier, we observed an increase transcription in these heterochromatic regions in PfHP1 mutants E-A, K-A and W-A when compared to the wild-type (Fig. 6A). Hi-C data of these mutant revealed that there is a weakened contact frequency at the heterochromatic regions (Fig. 6B and Supplementary Fig. 12 A,B). Also, there was an increase in the contact frequency within the euchromatic regions and between euchromatic and heterochromatic regions. Indicating that PfHP1 mutants lead to distinct changes in overall 3D genome organization.

**Figure 6:**
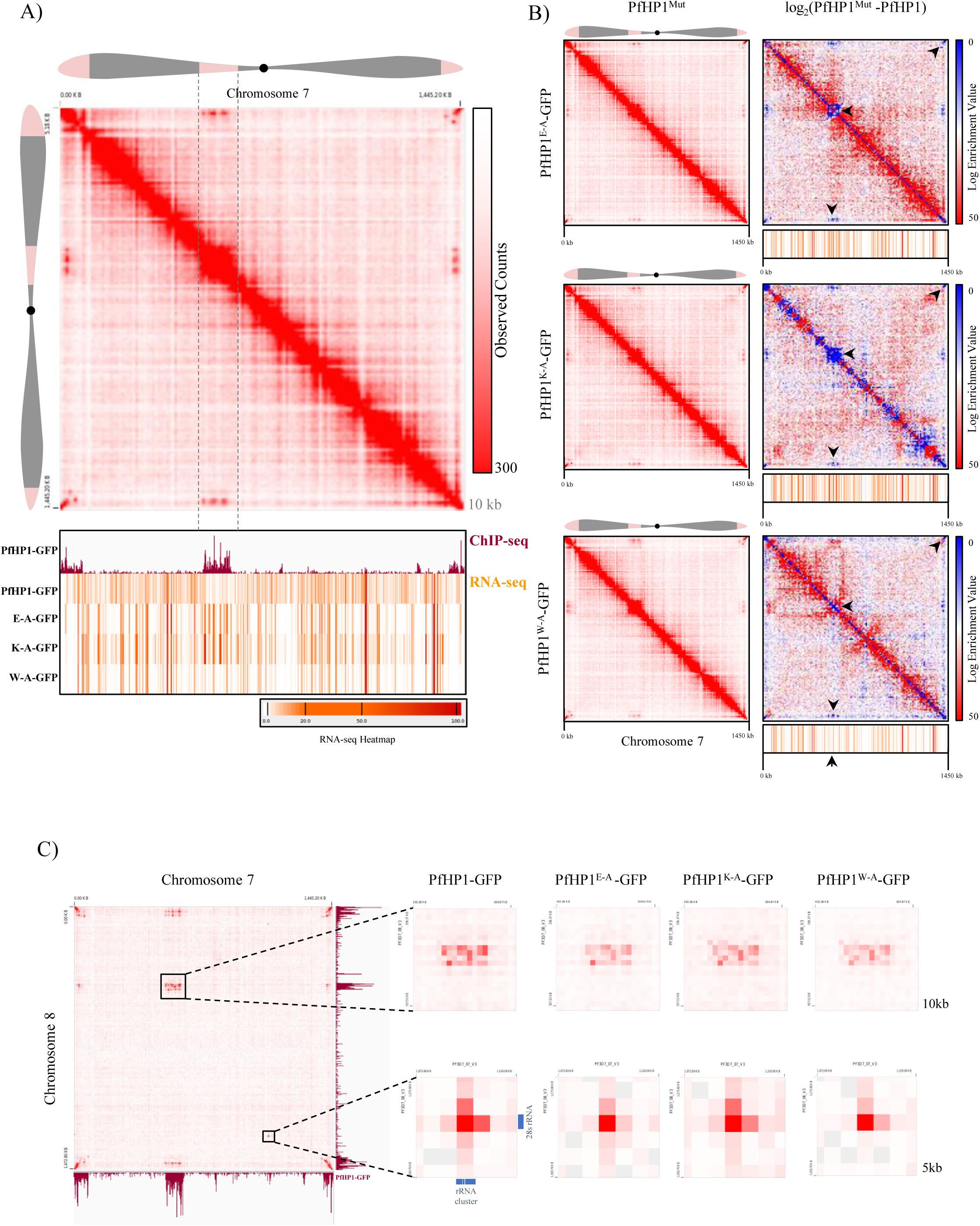
Hi-C contact map and features of GFP-tagged PfHP1 and its mutants. A) Hi-C interaction (visualized on Juicebox) within Chromosome 7 of PfHP1-GFP transgenic line (top) and comparison with PfHP1-GFP ChIP-seq peaks and the RNA-seq (for wild-type along with mutant lines E-A, K-A and W-A) visualized on IGV (bottom). The intensity of the red color indicates the frequency of contact. The RNA seq, for all three PfHP1 mutants and PfHP1-GFP control line, the corresponding bigwig file (used only replicate 1 for the visualization purpose) is visualized in the form of heatmap on IGV where orange denotes mid-range of number of transcripts and red indicates higher range of transcripts. The schematic of the chromosome 7 represents heterochromatic regions (pink color), euchromatic regions (gray color), and centromere (black dot in the middle). B) Intrachromosomal contacts of chromosome 7 are visualized on Juicebox for all the PfHP1 mutants. The contact frequencies are compared with PfHP1-GFP control line using the (Observed-Control) feature of juicebox. The loss of contacts between the *var* gene clusters are highlighted with black arrows. The HiC map is also compared with the RNA seq heatmap (at the bottom) for corresponding mutant lines. C) Inter-chromosomal contacts between chromosome 7 and 8 and a closer look at reduction in contact frequency across all the mutants of PfHP1(box on the top). Contacts between rRNA clusters are visualized at 1kb resolution and the contact is maintained even in PfHP1 mutants, shows that no-HP1 mediated clusters are not affected (box at the bottom).

Inter-chromosomal contacts between chromosomes were also disrupted specifically in the heterochromatic regions. The contact frequency of internal heterochromatic island between chromosome 7 and 8 is indicated in the Fig. 6C. All three mutations show loss of contacts in the region (Fig. 6C and Supplementary Fig. 13). However, a cluster of ribosomal genes did not show such an effect. Similarly, contact between the heterochromatic regions of chromosomes 4 and 7 decreased, while interactions between their centromeres became stronger in the mutants (Supplementary Fig. 12 C,D). These results indicate that PfHP1 mutations specifically affect both inter and intra chromosomal contacts of *var* genes and other CVGs. In conclusion, PfHP1 phase separation is an essential mechanism for the genome compartmentalization into euchromatin and heterochromatin.

## Discussion

Antigenic variation in *Plasmodium falciparum* is orchestrated by a multifaceted epigenetic program involving histone modifications, non-coding RNAs, and nuclear positioning (11–14,16,17,33,34,39,41,42,67–70). However, how these layers interact to enforce singular *var* gene expression remains poorly understood. To address this, we employed a multidisciplinary approach combining next-generation sequencing, gene editing, protein biochemistry, and single-molecule DNA assay.

We began with understanding the regulatory architecture of virulence gene expression, we examined the intronic regions of *var* genes, which emerged as hotspots for dynamic heterochromatin transitions from our ChIP-seq analysis (Fig. 1). The expansion and retraction of heterochromatin boundaries at these introns correlated with divergent transcription. This internal expansion and retraction of boundaries is consistent with the one of the four heterochromatin transitions identified by Michel-Todó et al., 2023 (71).The intronic regions also produced abundant sense transcripts, which we propose may: (i) poise *var* genes for activation, and/or (ii) limit heterochromatin spread, contributing to its maintenance rather than direct repression. The second scenario goes hand in hand with our observation later, where ncRNAs dissolve PfHP1 droplets in a dosage-dependent manner.

While heterochromatin is traditionally viewed as a repressive compartment, its role in *var* gene regulation appears more nuanced. Given the dynamic spatial and temporal changes in *var* gene expression during the intraerythrocytic developmental cycle (IDC), the underlying heterochromatin structure must be adaptable, dynamic and spatially separated. Our findings suggest that phase separation of PfHP1 provides a comprehensive mechanism that accommodates these aspects of gene regulation. PfHP1 forms dynamic, phase-separated condensates in the nucleus that likely facilitate the clustering and compartmentalization of *var* genes and associated regulatory proteins. These droplets are responsive to specific cues in concentration-dependent manner, such as ncRNAs and histone variant PfH2A.Z, suggesting a mechanism for tuning the heterochromatin states (Fig. 2). Given the requirement for precise temporal gene expression throughout the IDC, phase separation property may offer the speed and reversibility necessary for such tight regulation. Fluorescence recovery after photobleaching (FRAP) experiments further demonstrated that PfHP1 droplets are highly dynamic, supporting the hypothesis that these condensates allow the selective diffusion of macromolecules, while excluding components of active transcription machinery. We propose that, PfHP1 condensates selectively partitions perinuclear repressive centers that are composed of *var* genes, PfHP1, sirtuins and potential silencing factors from the RNA Pol II and other general transcription factors.

We also show the co-condensation of PfHP1 and parasite DNA in an AT-rich sequence-dependent manner (Fig. 3). While mammalian HP1α has been shown to form co-condensates with DNA, our findings provide the first evidence of DNA sequence-dependent co-condensation by PfHP1; marking the first such observation among all HP1 homologs. We propose a model where co-condensation of PfHP1 and AT-rich *var* intron/promoter sequences might be acting as nucleation centers (Supplementary Fig. 6F). This aligns with a recent study that has identified *var* promoters to be nucleation sites for *de novo* heterochromatin formation (72). We speculate the heterochromatin formation is a multi-step process where intermolecular interaction with DNA-PfHP1 helps in nucleation, and these condensates capture more DNA and start the compaction. While there could be additional DNA sequence determinants and DNA binding proteins involved in heterochromatin nucleation, it remains largely unstudied in the field. The multivalent interaction of PfHP1 with itself and H3K9me3 may help in the heterochromatin spread and droplet formation. Further fusion of multiple such droplets may lead to heterochromatin domain formation (Supplementary Fig. 6F).

We then extended our *in vitro* findings to the cellular context. ThePfHP1 phase separation mutants revealed defects in nuclear puncta formation, chromatin binding, and *var* gene silencing (Fig. 5). Notably, these mutants triggered aberrant activation of multiple *var* genes, reinforcing the critical role of PfHP1-driven compartmentalization in transcriptional repression. The L-D mutant showed most drastic effect on the heterochromatin gene expression; however, this mutant did not show defect in phase separation *in vitro*. While we speculate this to be a dimer mutant based on its homology to Swi6 DimerX mutant, the biochemical nature of this mutation needs to be seen in the future (56). Importantly, several of these phase separation mutants phenocopied the growth defects observed upon complete loss of PfHP1, highlighting the essential role of its phase separation in parasite viability (18,59). This phenotypic convergence strongly supports the notion that PfHP1-mediated compartmentalization is not merely supportive but central to maintaining heterochromatic repression and proper *var* gene regulation. The Hi-C data supports this notion by showing disruption of intra and inter-chromosomal heterochromatin contacts in the PfHP1 mutants.

Earlier it was shown that depletion of PfHP1 results in the disruption of repressive cluster. The CTCF, a major protein involved in genome boundary formation and organization in other eukaryotes is absent in *P. falciparum*, hence PfHP1 might be playing a central role in overall genome organization. We show that phase separation might be the fundamental mechanism for such a function (Fig. 6).

Based on these findings, we propose a speculative model for PfHP1 mediated *var* gene silencing (Fig. 7): PfHP1 forms phase separated condensates that compartmentalize *var* genes into repressive domains, physically segregating them from the transcriptional machinery. Intronic regions of *var* genes may serve as molecular “door handles” that remain outside the condensates, allowing regulated transcriptional access. Local increases in intronic transcription or ncRNA abundance may dissolve these condensates, thereby enabling *var* gene activation. However, the how the choice of *var* gene/s for intronic transcription needs to be seen. Our study introduces phase separation as a central mechanism in *Plasmodium* epigenetics, providing a new dimension to the regulation of clonally variant genes.

**Figure 7:**
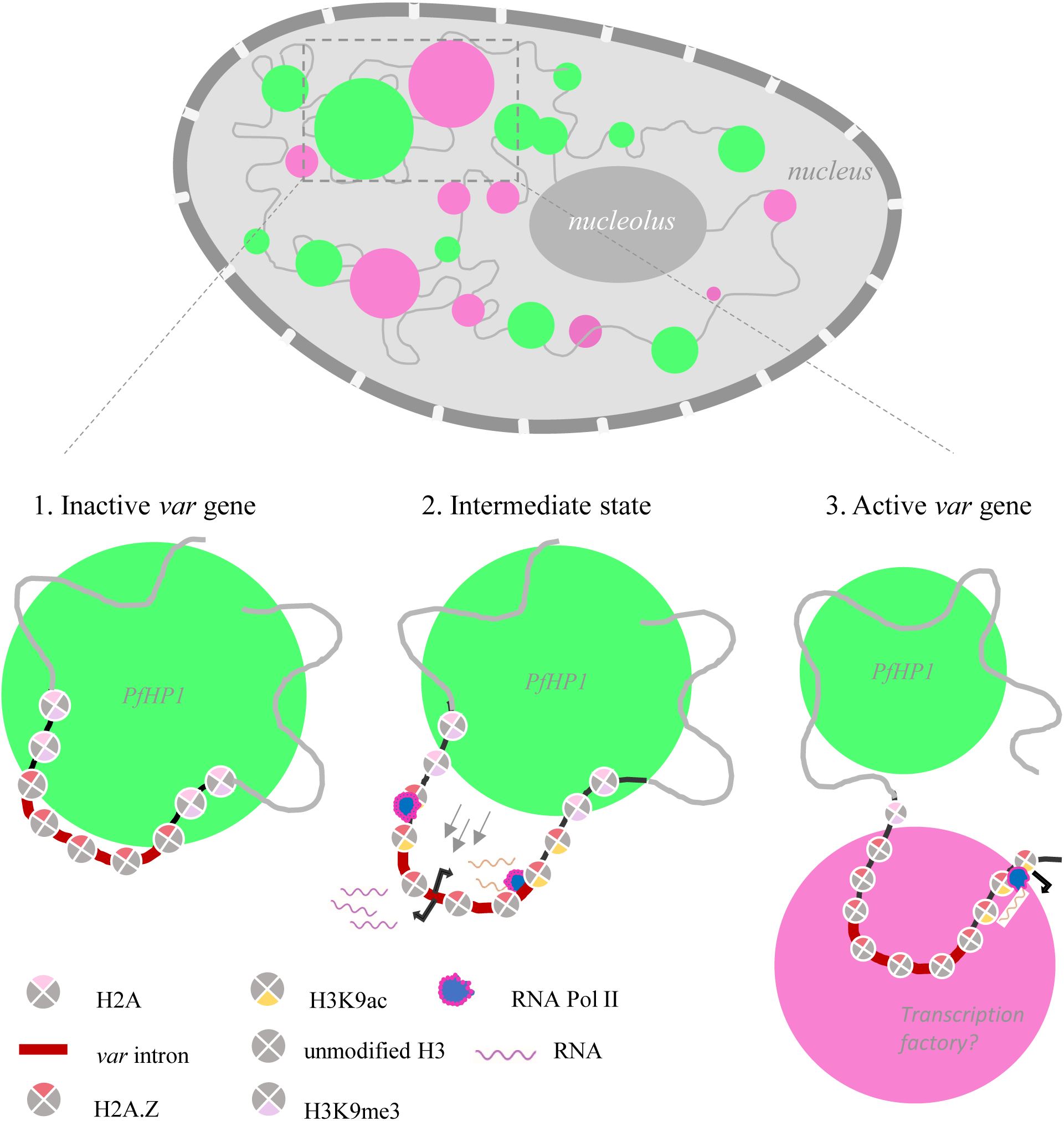
Model for PfHP1 phase separation-mediated regulation of *var* genes. A speculative model for the study representing the PfHP1 mediated *var* gene regulation via phase separation. In an inactive stage *var* genes are clustered together in a droplet of PfHP1 and the introns are looped out of the droplet. In an intermediate stage where the intron becomes more exposed and accessible to the transcription machinery and produces divergent transcripts. The active state is where the *var* gene re-localizes to a much more transcriptionally permissive environment and starts to produce full length transcripts. The local dissolution of droplets by *var* intronic ncRNAs/PfH2A.Z and other unknown nuclear components leads to coming out of the *var* gene from the droplets.

The biomolecular condensates formed by proteins is emerging as a new avenue for therapeutic targeting (26,28). This has already been extensively studied in diseases associated with protein phase separation, such as cancer and Alzheimer’s (73–75) but remains untapped in malaria. While challenges in achieving parasite-specific targeting remain, our findings on PfHP1 phase separation mutations may serve as a lead site for future drug targeting. In this light, this study on PfHP1 phase separation offers not only a conceptual advance in understanding the biology of *Plasmodium* but also a promising frontier in antimalarial strategy.

## Material and Methods

### Sequence analysis and alignment

The fasta sequence of HP1 homologs were obtained from NCBI database and the sequences were used to generate percent identity matrix, from sequence alignment algorithm of ClustalΩ (https://www.ebi.ac.uk/jdispatcher/msa/clustalo) (76). The percentage identity matrix was further represented as a heat map using Morpheus (Broad institute) (https://software.broadinstitute.org/morpheus/). For the prediction of IDRs PONDR (https://www.pondr.com/) was used (77). The propensity to phase separation was predicted using PSPredictor (http://www.pkumdl.cn:8000/PSPredictor/) (78).

### AT-content calculation of the gene

For the AT content calculation of multiple genes together, a sliding window frequency of 50 bp was scanned across the gene length, and the AT content was plotted using the Python package Matplotlib. For the calculation AT-content of individual genes, GC content calculator from biologics corp was used with a bin size of 30 bp (https://www.biologicscorp.com/tools/GCContent/). The gene sequences were retrieved from PlasmoDB (https://plasmodb.org/) and the promoter sequences were retrieved by selecting regions in IGV (integrative genomics viewer) against the *P. falciparum* genome.

### Cloning

Full length PfHP1 is cloned in the pET28a+ bacterial expression vector with His tag. The clones generated for the recombinant protein expression were, PfHP1 with GFP and His-tag in pET15b, PfHP1, PfH2A and PfH2A.Z with his tag in pET28a+ backbone. For the cloning of *var* intron into 18kbp plasmid the *var* intron (Pf3D7_0412400) was amplified from genomic DNA of 3D7 line (Primers; F-TGTGGGGTGAAAGCTGTAAAAAATATATTGTGGCGTATTT, R-CGGCAGAGTCTCTAGCTACAAAAAAATGAACACATATGT). Due to the extreme AT-richness we had to use reduced extension temperature (64°C) for the amplification.

### Site directed mutagenesis

The mutations generated were E6A/E7A/E8A, K17A/K18A/K19A/K20A, W29A, L254A, L254D in PfHP1-GFP (pET15b backbone) and PfHP1 (in pET28a+ backbone). In addition, Ser33 and Ser206 were mutated to cysteine in PfHP1 (pET-Ht backbone) for the fluorescent labelling purpose. The backbone plasmid was amplified with the primers with the mutations (see the list of primers is provided *Supplementary* Table 1). Pfu turbo (Agilent) enzyme was used in the PCR amplification. The PCR reaction was then subjected to DpnI (NEB) digestion (37°C, overnight) to remove the parent vector. From the reaction 2μl was transformed into DH5α and plated on a Luria agar plate with appropriate antibiotic selection. The clones were screened and sequence verified for the mutation.

### Bacterial expression and purification of PfHP1 recombinant proteins

The vector was transformed in Bl21 (DE3) *Escherichia coli.* The culture was induced for protein expression using 0.5 mM IPTG (isopropyl-1-thio-β-d-galactopyranoside). The induction was done overnight at 18°C. The pellet was spun down and stored at -80°C until the protein purification. The cell pellet was resuspended in sonication buffer (HEPES pH 8.0: 20mM, KCl: 300mM, imidazole: 10mM, Triton X100: 0.5%, PMSF: 1mM, Glycerol: 10%, Lysozyme: 1 mg/ml,β-mercaptoethanol (β-me): 2mM; 40 ml sonication buffer for the pellet from 1L culture) and sonicated for 40 minutes at 60% amplitude, 2 sec ON and 5 sec OFF cycles in a probe sonicator (Vibra-cell, Sonics and Material Inc.). The lysate was spun at 12000 rpm, 4°C, 20 min. The supernatant was allowed to bind to Ni-NTA beads for 2 hours in rotation at 4°C. The unbound lysate was removed by gravity flow. The beads were washed with 5 CV (column volume) of wash buffer I (pH 8.0 HEPES: 20mM, KCl: 1M, Glycerol: 10%, Imidazole: 10mM, β-me: 2mM), wash buffer II (pH 8.0 HEPES: 20mM, KCl: 300mM, Glycerol: 10%, Imidazole: 30mM, β-me: 2mM) and wash buffer III (pH 8.0 HEPES: 20mM, KCl: 300mM, Glycerol: 10%, Imidazole: 50mM, β-me: 2mM). The PfHP1 was eluted in 1 CV of elution buffer (pH 8.0 HEPES: 20mM, KCl: 300mM, glycerol: 10%, Imidazole: 500mM, β-me: 2mM) a total of 5 elutions were taken. The protein was estimated using spectrophotometric measurement or a nanodrop. The protein quality was assessed by running on SDS-PAGE gel (Supplementary Fig. 14A). The protein was stored at 4°C for short term and -80°C for long term storage. The same protocol was followed for PfHP1-GFP (and all the mutant PfHP1 made in the backbone) except we did not add β-mercaptoethanol in the buffers. After the purification, proteins were concentrated using amicon 10kDa and 30kDa membrane filters (millipore) for PfHP1 and PfHP1-GFP respectively. The concentrated proteins were desalted/buffer exchanged (20mM HEPES pH 7.4, 300mM KCl, 5% glycerol, 2mM EDTA) using PD-10 column (Cytiva). The proteins were further concentrated if needed using amicon centrifugal filters. The protein quality was assessed by running on SDS-PAGE gel (Supplementary Fig. 14D). The proteins were quantified using absorbance at 280 nm using a spectrophotometer. The freeze-thaw cycles were avoided and the experiments were performed with freshly desalted proteins.

### Bacterial expression and purification of histone recombinant proteins

The expression conditions used for PfH2A and PfH2A.Z were the same as that of PfHP1. The cell pellet was resuspended in sonication buffer (50mM Na_3_PO_4_ pH 8.0, 500mM NaCl, 20mM imidazole,10% glycerol, 3mM β-ME, 1mM PMSF, 0.5% TritonX100, 0.5mg/ml lysozyme; 40 ml sonication buffer for the pellet from 1L culture) and sonicated for 40 minutes at 60% amplitude, 2 sec ON and 5 sec OFF cycles in a probe sonicator (Vibra-cell, Sonics and Material Inc.). The lysate was spun at 12000 rpm, 4°C, 20 min. The supernatant was allowed to bind to Ni-NTA beads for 2 hours in rotation at 4°C. The unbound lysate was removed by gravity flow. The beads were washed with 5 CV (column volume) of wash buffer (40mM Imidazole, 50mM Na_3_PO_4_ pH 8.0, 500mM NaCl, 3mM β-ME, 0.1mM PMSF). The Proteins were eluted in 1 CV of elution buffer (300mM Imidazole, 50mM Na_3_PO_4_, 500mM NaCl, 3mM β-ME, 0.1mM PMSF) for a total of 5 elutions. After the purification, proteins were concentrated using Amicon 10kDa membrane filters (millipore). The concentrated proteins were desalted/buffer exchanged (20mM HEPES pH 7.4, 300mM KCl, 5% glycerol, 2mM EDTA) using PD-10 column (Cytiva). The proteins were further concentrated if needed using Amicon centrifugal filters. The purity of the protein was checked using SDS-PAGE. And quantified using absorbance at 280 nm in a spectrophotometer or a nanodrop.

### Antibody generation

The purified PfHP1 protein was used as an antigen for the antibody generation. The antibodies were raised in The National Facility for Gene Function in Health and Disease, IISER Pune. The New Zealand White rabbit was immunized with 300 µg of protein with complete Freund’s adjuvant (CFA). The antigen-adjuvant emulsion was made by mixing equal volumes of the protein and CFA using a three-way syringe. Before the first injection collected pre-immune sera. For the subsequent booster shots 200 µg protein was injected along with Incomplete Freund’s adjuvant. After the first booster shot the bleed was collected every 14 days. The immune sera was separated from the whole blood by centrifugation. The antibody was validated by Western blot with *P. falciparum* nuclear lysate (Supplementary Fig. 14B,C).

### Western blot for antibody validation

For the antibody validation *P. falciparum* nuclear lysate was prepared following the given protocol. Briefly the parasites were enriched by saponin (0.015%) lysis of RBCs. The pellet was first lysed by a cytoplasmic buffer (10mM HEPES pH-7.9, 10 mM KCl, 0.1 mM EDTA, 0.1 mM EGTA, 0.65% NP40, 1 mM DTT, 2X PMSF) and then by a high salt nuclear extraction buffer (20 mM HEPES pH 7.9, 400 mM NaCl, 1 mM EDTA, 1 mM EGTA and 1 mM DTT). The nuclear extract (25 µg) was used for western blotting. After the transfer, PVDF was blocked using 5% milk and probed with primary antibody (in-house PfHP1 antibody in 1:200 dilution) overnight. Next day, probed with anti-rabbit HRP (Horseradish peroxidase) antibody in 1:5000 dilution. Gave three washes with TBST (1X Tris-buffered saline, 0.1% Tween 20) and developed the blot using Clarity max ECL substrate (Bio Rad).

### Fluorescence labelling of the protein

S33C and S206C mutants of PfHP1 were labeled with fluorophores under denaturing conditions at pH 7.4. The proteins were labelled by Alexa Fluor-488 dye (C5-maleimide), in a 2:1 molar ratio (dye:protein). The reaction mixtures were stirred for 2-3 hours in the dark at room temperature. After the labelling the excess dye was removed by PD-10 columns (20 mM HEPES pH 7.4, 300mM KCl, 5% glycerol, 2mM EDTA), and concentrated using a 10 kDa MWCO Amicon membrane filter. The concentration of the labeled protein was estimated using absorbance at 280 nm and 495 nm (Alexa Fluor 488 C5-maleimide ε495 = 72,000 M-1cm-1). 1% of the labelled protein was added to the unlabelled PfHP1 for phase separation experiments.

### YOYO-1 labelling of the DNA

The 601 Widom sequence was amplified from a pGEM 601 vector (addgene #26656) using specific primers (*Supplementary* Table 2). The PfDNA (*P. falciparum* DNA) is an 800 bp stretch amplified from the *P. falciparum* genome (primers are given in the *Supplementary* Table 2). The 601 Widom DNA was labelled with YOYO-1 in a ratio of one molecule of the dye per 10bp of DNA.

### In vitro transcription of RNA

The template for the IVT is PCR amplified. The GC-rich RNA was coded from the RUF6 gene in the *P. falciparum*, the primer used contained overhangs with T7 promoter and terminator for the IVT reaction. The template for AU-rich RNA was difficult to amplify because of the extreme AT-richness. Overhanging oligos were used to assemble the template for the AU-rich RNA (oligos given in the *Supplementary* Table 3). The PCR products were cleaned up using Phenol:chloroform:isoamyl alcohol method. The IVT reaction was set up according to the stanard protocol given for Invitrogen T7 polymerase. The reaction was kept overnight at 37°C in a PCR machine. Next day DNaseI (Promega) treatment at 65°C for 10 minutes was performed. The reaction was stopped by adding DNaseI stop solution. The IVT RNA was cleaned up using precipitation by 2.5V 100% ethanol and 0.1V ammonium acetate. To ensure that the obtained product is RNA and not DNA, RNaseA (Himedia) digestion was performed. The purity of the RNA was inspected by running the samples with or without RNaseA on a 1.3% agarose gel. The quantification was done by a nanodrop instrument.

### Droplet formation assay

All phase separation were adapted from Rai et al., 2023 (79). Briefly the protocol is, before the droplet assay proteins were spun at a high speed to remove any possible debris or precipitate. In a clean PCR tube, mix 10μl of buffer (20mM HEPES, pH 7.4) and 10 μl of PfHP1-GFP/mutants (30μM stock) or PfHP1 (100μM stock), this is the basic droplet assay reaction mix. Gently pipette and avoid bubbles. Observed under a microscope by putting 5μl of the reaction on a coverslip of 1.5 mm thickness. The microscopes used were ZEISS LSM 980 Elyra 7 or ZEISS LSM 710 confocal microscope. The proteins were imaged under 488 nm laser, 63x oil objective. The other conditions tested were different temperatures and salt concentrations. Also effect of 601 Widom DNA, PfDNA, RNA (IVT AT-rich and GC-rich ncRNAs), PfH2A, PfH2A.Z, and PEG8000 were tested. The test component was added to the pre-formed PfHP1 droplets. The amount of protein and other components used are mentioned along the figures. For the droplet area quantification Fiji (Image J) software was used. The images were converted to gray scale (8 bit) first and changed the threshold setting to ‘intermodes’. The image was then analysed using the ‘analyse particles’ option to obtain droplet area, number and mean intensity.

### Fluorescence recovery after photobleaching (FRAP) for phase separated droplets

FRAP experiments were performed on ZEISS LSM 980 Elyra 7 super-resolution microscope. For all FRAP experiments, GFP tagged PfHP1 was used. FRAP measurements were performed for at least three independent samples. A region of interest (ROI) of 1μm diameter was defined within a selected droplet, and a same size ROI was defined outside as background. The ROI within the droplet was bleached using a 488 nm laser; five frames were taken before the bleaching and recovery was followed for 70 seconds. The recovery of the bleached spots was recorded using the in-built ZEN blue 3.2 (ZEISS) software. The five different data sets were background corrected using the outside ROI, normalized with standard deviation, and plotted using Origin 2020b.

### Saturation concentration estimation

Prepared 500μl droplet assay reaction in a 1.5 ml tube with 15μM PfHP1 protein in 20mM HEPES pH 7.4, 150mM KCl, 2.5% glycerol and 1mM EDTA buffer. Incubated the reaction at room temperature for 5 minutes and spin at 16000 rpm, 25°C for 35 minutes. Estimated the volume of supernatant after the spin and back calculate the volume of pellet or dense phase. Measured the absorbance at 280 nm of the light phase/supernatant, this shall give you the saturation concentration or C_sat_. Similarly measured the dense phase absorbance by resuspending the pellet in 80μl urea 8M (in 1X TBS). The concentration was calculated from the absorbance value. The experiment was performed in triplicates.

### Turbidity assay

For the turbidity assay with DNA, 20μl reaction was prepared with varying concentration of PfDNA (1, 2, 5, 10, 20, 40, 80, 100 ng/μl) and 15μM PfHP1 in 20mM HEPES pH 7.4, 150mM KCl, 2.5% glycerol and 1mM EDTA buffer. The reaction was incubated at room temperature for 5 minutes and measured absorbance at 340 nm. All the experiments were performed with triplicates.

### EMSA (Electrophoretic mobility shift assay)

The 1.3% agarose gel is made in 1X Sodium borate buffer (Stock: 20X SB buffer, pH 8.0, 8g NaOH and 48g Boric acid in 1L water). The PfDNA was made by amplifying the *var* intron from a 19kbp vector using the same primers used for cloning. The DNA was mixed in 20 ng/μl concentration with increasing concentration of PfHP1 (0nM, 125nM, 250nM, 500nM, 1μM, 2μM, 4μM and 10μM). The reaction was incubated for 30 minutes at room temperature and then run on the gel at 150V for 1 hour. Visualized the gel using EtBr.

### Biotinylation of the 18kbp DNA and 19kbp DNA with var intron

The 18 kbp (control) and 19 kbp (with *var* intron) vectors were digested with NotI and XhoI (NEB) and similarly 500 bp biotin-tagged PCR fragments (biotin handles) digested with the same enzymes. For Cy5-labeling, Aminoallyl-dUTP-Cy5 (Jena Bioscience) was incorporated in one of the biotin handles and then digested this with NotI enzyme and other handle with XhoI enzyme. The ligation reaction was set-up using T4 DNA Ligase and ran the ligated product on 0.8% agarose gel to separate ligated product from non-ligated product. The resultant length of the DNA was 19 or 18kb with or without Biotin-dUTP-Cy5 at one end and Biotin-dUTP at another end. The Cy5 labeling helps in determining the orientation of the DNA molecule.

### Preparation of PEG passivated slides

Slides were prepared following the protocol from Satheesh et al., 2025 (80). Briefly, slides were drilled, cleaned via sequential sonication (detergent, Milli-Q, acetone, KOH, and Piranha solution), and then silanized using APTES/acetic acid/methanol. PEG passivation was achieved using a 1:10 mix of biotin-PEG-SVA and mPEG-SVA (Laysan Bio) in freshly prepared 0.1 M K SO (pH 9–9.5). After overnight incubation, slides were rinsed and subjected to a second PEGylation using MS(PEG) (Thermo Scientific).

### Single molecule DNA tethering experiment

Microfluidic flow cells were constructed using double-sided tape between quartz slides and coverslips. Internal surfaces were passivated with 50:1 mPEG/biotin-PEG (20% w/v in 100 mM K SO). Channels were treated sequentially with 10% BSA, 0.1% Tween-20, and T50 buffer (50 mM Tris pH7.5, 50 mM NaCl). Neutravidin (0.2 mg/ml) was incubated to enable biotin-based DNA tethering. Biotinylated DNA (∼10-50pM) was introduced at 6–15 µl/min using a syringe pump. The tethered DNA was visualized under the microscope using DNA intercalating dye 50nM Sytox Orange (Thermofisher Scientific S11368, excitation 561 nm). Proteins (PfHP1-eGFP or mutants) were added at varying concentrations for specific experiments. Imaging was performed on a TIRF microscope (Nikon Ti2 Eclipse) using a 100× oil immersion objective (NA 1.49, IISc Bangalore), under total internal reflection conditions.

### Data analysis of single molecule experiment

Fluorescence images were acquired using NIS Elements and converted to tiff format for analysis in custom MATLAB software. Normalized intensity profiles were concatenated over time to generate kymographs, which were used to estimate the base pairs of DNA compacted regions using the following formula;

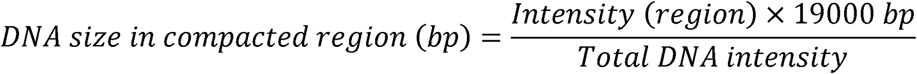

To analyze the 19 kb DNA molecules containing a central *var* intron AT-rich region, individual DNA strands were first captured using Fiji ImageJ. From these images, a time-averaged projection was created by averaging the fluorescence intensity across 200 frames. The end-to-end distances of the DNA molecules were then normalized on a scale from 0 to 1, allowing the precise positioning of the condensate along the DNA strand to be determined. The resulting data was further processed through interpolation and averaging to produce the plot shown in the results section. The puncta number was calculated from the maximum intensity projections made using Fiji Image J. The data was plotted using graph pad Prism 10. The significance and P value were calculated by non-parametric t-test.

### Parasite culture

The *P. falciparum* line 3D7 was maintained in 2% hematocrit in RPMI 1640 10.4g/liter, HEPES 25 mM, AlbuMAX II 0.5% (wt/vol), hypoxanthine 370μM, gentamicin 12.5μg/ml, sodium bicarbonate 1.77 mM (just before use by adding 3 ml of 5% (wt/vol) sodium bicarbonate to 100 ml of medium). The parasites were maintained at 37 °C in an atmosphere with 5% carbon dioxide, 5% oxygen and rest nitrogen. The parasites for the experiments were synchronized by 5% sorbitol and 63% percoll gradient cyclically to achieve a tight synchrony.

### Transfection constructs: Ectopic expression lines expressing PfHP1 mutants

Inducible expression lines of wild-type PfHP1 and mutants were generated by targeting the *cg6* locus (PF3D7_0709200, a dispensable locus) of 3D7/DiCre (61). The guide RNA against the *cg6* locus was cloned in the pBF_gC plasmid (66,81). The transgenics are made by co-transfecting 50μg each of pBF_gC_cg6 and donor plasmid pD_gC_cg6_pfhp1_gfp_dd_glmS. The mutant PfHP1 proteins are under calmodulin (cam) promoter and conditionally regulated by glmS ribozyme and FKBP-DD (destabilising domain). In the presence of Shield-1, the DD domain is stabilised and allows mutant PfHP1-GFP expression, while Shield-1 removal leads to proteasomal protein degradation (82). The glmS ribozyme cleaves the ectopically expressed PfHP1-GFP mRNA upon the treatment with glucosamine (83).

### Transfection constructs Conditional expression of pfhp1 mutants at the endogenous pfhp1 locus

The donor plasmid for the CRISPR was cloned in the pUC backbone using Gibson assembly cloning technique. The design of the construct is given in Fig. 5A. The guide RNA targeting the *pfhp1* loci was cloned in a pHF_gC backbone. The CRISPR strategy was based on a self-cleaving ribozyme that will simultaneously give rise to two gRNAs once transcribed inside the cell. The two gRNAs (gRNA 1 and gRNA 2) will target the cas9 to cut and homogeneously recombine at the beginning and end of the gene. The gRNAs were designed using CRISPOR (https://crispor.gi.ucsc.edu/) online tool (84). The stable knock-in lines will code for the wild type PfHP1 with the mScarlette-I tag. Upon rapamycin treatment, the 3D7/1G5/DiCre lines will start dimerising Cre recombinase, which will recombine loxP sites and produce GFP-tagged mutant PfHP1 protein. 50μg of the plasmids were precipitated with 0.1V sodium acetate and 2.5V absolute ethanol at room temperature for 10 minutes. Washed with 70% ethanol. Resuspended it in 1X TE (25 μl).

### Transfection

The transgenic lines were generated in 3D7/1G5/DiCre background (61). The transfection was done by following the previously published protocol (66). Before transfection, sorbitol synchronizes the 3D7/DiCre line and enriched early ring stages. Taken 250 μl of culture pellet with ∼5% parasitemia and wash (600g, 4 min) the pellet with 2 ml cytomix (120mM KCl, 0.15mM CaCl_2_, 2mM EGTA, 5mM MgCl_2_, 10mM K_2_HPO_4_/KH_2_PO_4_ 25mM HEPES, pH 7.6).

Removed the supernatant and added 400μl cytomix and 25μl each of the plasmids (50μg) that need to be co-transfected. Resuspended with pipetting and incubate at room temperature for 5-10 minutes. Transferred the mix to a cuvette with 1ml pipette without any bubbles. Immediately gave electric pulse in a Biorad electroporator with the following program 310V/960µF. Immediately after the pulse, the cells are transferred to a 100 mm dish with 10 ml media and 400 μl of fresh blood. Kept it in an incubator in 37°C temperature and 92% N_2_/4% CO_2_/3% O_2_. Changed media in the next 24hrs and start drug selection with the appropriate drug (WR 20uM stock: used 5nM, Bsd 10 mg/ml: used 5μg/ml). Add WR drug every day until 7-8 days, or till all the parasites are cleared. Add Bsd drug for every 9-10 days. All the lines were maintained in 5% hematocrit in RPMI 1640 medium supplemented with 25 mM HEPES, 370 µM hypoxanthine, 24 mM sodium bicarbonate, and 0.5% AlbuMAX II. Also, to reduce background conversion into sexual stages (85), the complete medium was complimented with 2mM Choline Chloride.

### Glucosamine and Shield-I treatment

The ectopic mutant expressing lines were continuously maintained in the 2.5mM glucosamine from the day of transfection. For the harvesting and expression of the mutants the glucosamine was removed and 1.25 μM Sheild-1 was added to the culture. The next day the mutants were used for the experiments.

### Rapamycin treatment

Double synchronized the PfHP1 mutant lines in the 3D7/1G5/DiCre background using 5% sorbitol. The early ring parasites treated for 4 hours with 100nM rapamycin. The parasites were harvested after 48 hours in next cycle. The PfHP1-GFP wild-type was used as a control in all experiments.

### Parasite harvesting

Spun down the culture at 650 g, 5min. To the culture pellet added 0.015% saponin and spun at 2000 rpm, 5 min. Washed with 0.015% saponin for 1-2 times Washed with 1x PBS 1-2 times. If harvesting for RNA isolation after this step, added trizol and thoroughly resuspended the pellet, and stored at -80°C. If harvesting for protein, lysed the cell pellet with appropriate lysis buffer and stored the cell lysate at -80°C.

For ChIP: Ring stage (∼10 hours post infection, hpi) parasites were double sorbitol synchronized by 5% sorbitol. Spun down the culture at 650 g, 5min. Used at least 500ul culture pellet with 10% parasitemia per ChIP sample for ring stage ChIP. After spinning down, removed half of the media and add 1% formaldehyde (16% stock Formaldehyde methanol free, Thermofisher scientific) dropwise. Kept on a rocker at room temperature for 10 minutes, 15 rpm. For quenching, added 150mM glycine (2M stock). Kept on a rocker at room temperature for 10 minutes, 15 rpm. Spun at 6000 rpm, 4 °C, 5 minutes. Washed with chilled 1X PBS and spin the pellet at the same speed for 5 minutes (just reverse the pellet direction so that the pellet travels through the 1X PBS. Repeated the previous step for a total of 3 washes. Stored the pellet at - 80°C.

For RNA: Ring stage (∼10 hours post infection, hpi) parasites were double sorbitol synchronized by 5% sorbitol. The samples were spun down and harvested using 0.015% saponin. The cell pellet was resuspended in Trizol and snap frozen and stored -80^0^C until further proceeding.

### PCR validation

The genomic DNA was isolated from the wildtype and mutant lines using QIAGEN blood easy kit (Primers are given in *Supplementary* Table 4). PCR was set up using the primers upstream and downstream the insertions. NEB Q5 DNA polymerase was used for the amplification. The amplification products were run on 1% agarose gel and stained with EtBr for the visualization.

### Western blot validation of lines

The saponin harvested parasites were lysed in 8M Urea/10% SDS buffer. The supernatant after lysis was taken and quantified using Qubit Protein assay kit (High sensitivity or broad range for quantification). And ∼25μg was used for Western blot gel run. PVDF membrane (Bio Rad) was used for western transfer and 5% skimmed milk was used for blocking. Primary antibodies used was α-CD (1:5000) (18).

### Live cell fluorescence imaging and FRAP assay

The parasites were labelled with Hoechst/DAPI (Invitrogen) for 30 min at 37°C. Cells were washed with 1X PBS and mounted on a slide with vectashield/slow fade antifade (Invitrogen). The images were taken at Leica DM6. The brightfield images are taken in Leica DM750 microscope. For the FRAP assay cell imaging dish 145uM, 35 x 10 mm (Eppendorf) was coated by poly-D-Lysine (Gibco) for 1 hr and then washed off with autoclaved MilliQ water and prepared for the live imaging of *Plasmodium falciparum* infected RBCs. The imaging was performed using Zeiss Multiphoton confocal microscope. Puncta of HP1-GFP were photobleached for less than 1 second with 100% laser power. For analysis, mean fluorescence intensity was measured using Fiji (ImageJ). The puncta showed dynamic movement in all three axes; thus, quantitative analysis was technically limiting. Although visually, the fluorescence recovery of around 80% occurs within one minute. The signal recovery and puncta were manually tracked and measured. The normalization was done according to this formula –

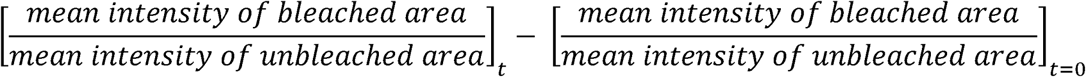

### Chromatin immunoprecipitation (ChIP)

The ChIP was done by following the previously published protocols (86,87). To the harvested *P. falciparum* parasites, added 6 times of swelling buffer (25 mM Tris pH 7.9, 1.5 mM MgCl2, 10mM KCl, 0.1% NP40, 1mM DTT, 0.5 mM PMSF, 1x PIC (Roche)). Kept on ice for 10 minutes and Dounced 100 times with a loose piston for the ring stage parasites. Spun at 5000 rpm for 5 minutes at 4^0^C. Discarded the supernatant and added the sonication buffer (50 mM Tris-Cl pH7.9, 140 mM NaCl, 1mM EDTA, 1% Triton X-100, 0.1% Sodium deoxycholate, 1.0% SDS, 0.5 mM PMSF, 1x PIC (Roche)) 8 times to the packed pellet. Sonication conditions for covaris M220 were; average incident power: 10.1 W, peak incident power: 75.0 W, Duty factor: 13.4%, cycle/burst: 200, Duration: 720 seconds. After the sonication, spin at 13000 rpm, 15 minutes at 4^0^C. Took 20-25μg of chromatin (for *P. falciparum*, samples can go as low as 10-20μg if the samples are limited). Made the volume up to 1.0 ml with a chromatin dilution buffer (0.01% SDS, 1.1% Triton X-100,1.2 mM EDTA, 16.7mM Tris-Cl pH 8.0, 167 mM NaCl). The pre-clearing step is optional if using protein A dynabead (Invitrogen). For the pre-clearing, added 10μl beads to the chromatin lysate for 1 hour, 4^0^C, and 10 rpm end-to-end rotation. Separated the supernatant using a magnetic stand and added the specific antibody. Primary antibodies used were α-GFP (Abcam, ab290), α-H3K9me3 (Invitrogen, PA5-31910), α-H3K9ac (Millipore, 06-942), α-H3K79me2 (Sigma, 04-835), α-H4K8ac (Sigma, 07-328), α -RNA Pol II Serine-2-phosphorylation antibody (Abcam, ab5095), α-PfHP1 (in-house), α-PfSir2A (in-house), α-H3K36me3 (Invitrogen, MA524687). Incubated overnight 4^0^C and 10 rpm. Next morning, added 40 μl of saturated beads to the lysate. To prepare saturated beads, incubated the beads with 0.4 mg/ml tRNA and 0.5 mg/ml BSA for 3 hrs, 4^0^C, and 10 rpm. The lysate was incubated with the beads for 4 hrs, 4^0^C, and 10 rpm. Separated the beads by a magnetic stand and washed thrice with the chilled low salt buffer (0.1% SDS, 1% Triton X-100, 2 mM EDTA, 20 mM Tris-Cl pH 8.0, 150 mM NaCl), high salt buffer (0.1% SDS, 1% Triton X-100, 2 mM EDTA, 20 mM Tris-Cl pH 8.0, 500 mM NaCl), LiCl buffer (0.25 M LiCl, 1% NP40, 1% Sodium deoxycholate, 1 mM EDTA, 10 mM Tris-Cl pH 8.0) and TE buffer (10 mM Tris-Cl pH 7.5, 1 mM EDTA). Incubated for 10 minutes with each buffer at 4^0^C and 10 rpm. Finally, eluted the DNA using 150μl elution buffer (10 mM Tris-Cl pH 7.5, 1 mM EDTA) at 37^0^C for 5 minutes. Repeat the step once more to get a total of 300 μl elute. The de-cross linking was performed, followed by DNA extraction. Estimated the DNA amount with Qubit DNA high-sensitivity kit (Invitrogen). Proceeded with the library preparation.

### ChIP library preparation and sequencing

The sequencing library was prepared using the NEB Ultra II DNA kit (E7645), following the manufacturer’s instructions. The prepared library was estimated for the average size using Agilent bioanalyzer or tapestation and for the amount using Qubit fluorometer. A final amount of 1.3 pM library was loaded from the multiplexed pool of all libraries. Paired end sequencing was performed using Illumina 300 cycle mid output kit. The NextSeq 550 (IISER Pune, NGS facility) was used for the sequencing.

### RNA isolation

The RNA isolation was done on the day of library preparation to minimise freeze thaw cycles. The samples were thawed and added 0.2 volumes of chloroform and mixed thoroughly by inverting. Incubate at room temperature for 15 minutes, followed by a spin at 12000 rpm, 15 minutes, 4^0^C. Separate the top layer and add 0.8 volume of isopropanol (may add glycogen during this step to increase the visibility of the pellet). Invert mix and incubate at room temperature for 30 minutes. Spin down at 12000 rpm, 15 minutes, 4^0^C. Observe a white pellet and wash the pellet with 70% ethanol. Dry the pellet at room temperature to evaporate and residual ethanol. Resuspend the pellet in 20 μl nuclease free water. The RNA quality was assessed using RNA high sensitivity nano chip for Agilent bioanalyzer or tapestation and the RNA was quantified using Qubit RNA high sensitivity kit.

### RNA library preparation and sequencing

The isolated RNA was subjected to polyA enrichment using NEB polyA mRNA magnetic isolation module (E7490). The library preparation was done using the NEB Ultra II RNA kit (E7770). And for the DiCre line samples NEBNext Ultra™ II Directional RNA Library Prep with Sample Purification Beads (E7765) was used for the library preparation. The library quality was tested using a DNA high sensitivity kit in Agilent bioanalyzer. The library amount was estimated using Qubit DNA high sensitivity kit. A final pool of 1.7 pM library was loaded on a 75 cycle High output illumina sequencing kit. The single end or paired end sequencing was performed using NextSeq 550 from Illumina at IISER Pune NGS (Next generation sequencing) facility.

### ChIP sequencing data analysis

Trimming and quality control (QC) of the raw data was done using Trim Galore. The trimmed data was aligned against the *P. falciparum* reference genome (PlasmoDB_v53) using Bowtie2. The aligned files were sorted using samtools which gave bam output files. The peaks were called using MACS2, using the input files as control. The normalized bedgraph files were loaded on IGV (Integrative Genomes Browser) to visualize the enrichment pattern. For the mean tag density and gene body profile seqMINER was used. The heatmaps were generated using Morpheus, Broad Institute.

### RNA sequencing data analysis

Trimming and QC of the raw data was done using trim galore. The trimmed data was aligned against the *P. falciparum* reference genome (PlasmoDB_v53) using HISAT2. The aligned files were sorted using samtools which gave bam output files. The tpm (transcripts per million) value for genes were obtained from StringTie analysis. The mean tag density for the heat maps were obtained from seqMINER software. The heatmaps were generated using Morpheus, Broad Institute. For the directional RNA sequencing analysis deeptools were used for generating bigwig files for the + strand and -strand. The RNA sequencing data was normalized using TPM (transcript per million) option. The heatmap visualization of the bigwig files were done using deeptools plotHeatmap. The differential enrichment analysis for the data was done using Deseq2. The wild-type PfHP1-GFP was used as a control for each mutant line. The heatmaps were generated using Morpheus, Broad Institute.

### Hi-C sequencing

50 ml culture of double-synchronized ring stage parasites at 3% hematocrit and around 10% parasitemia were crosslinked and prepared for Hi-C as instructed in Arima-HiC kit. An extra step of RBC lysis with 0.06% saponin was carried out before aliquoting and snap freezing. Hi-C data was generated using the Arima-HiC kit and Arima Library Prep Module according to the manufacturer’s protocols. The libraries were sequenced in a 75-bp paired-end run on the NextSeq 550 (Illumina) platform.

### Hi-C data analysis

Paired-end HiC library reads were processed (i.e., mapping, filtering, and pairing, and normalization) using Juicer 1.22.01 and the *P. falciparum* 3D7 genome (PlasmoDB v66) with mapping at multiple resolutions 50,25,10,5 and 1 kb resolution as the restriction sites (DpnII and HinfI) provide a sub 1kb resolution for *Plasmodium falciparum* genome (Supplementary data). The HiC contact heatmaps were visualized using Juicebox 1.11.08.

### Data access

PfH2A and PfH2A.Z ChIP sequencing data reanalyzed from Bartfai et al. 2010 (GEO accession number: GSE23787). The ATAC sequencing and stranded RNA sequencing data was reanalyzed from Toenhake et al. 2018 (GEO accession number: GSE104075). ChIP-sequencing, gene expression data (RNA sequencing) and HiC sequencing data are deposited in NCBI Gene Expression Omnibus (GEO) under the accession numbers GSE299944, GSE300206 and GSE300368 respectively.

### Ethics Statement

This study does not involve human participants. Human RBCs used in this study were obtained from the KEM Blood Bank (Pune, India) as blood from anonymized donors.

## Acknowledgements

We are thankful to Bhagyashree Deshmukh for the help with initial parasite growth and harvesting, Prof. Geeta Narlikar and Dr. Gayathri Pananghat for their valuable suggestions that contributed to the development of this project. The parasite line 3D7/1G5/DiCre used for PfHP1 mutant line generation was obtained from Michael J. Blackman. We acknowledge the use of BioRender (BioRender.com) for some of the illustrations and OpenAI’s assistance in language refinement in this manuscript. DVM is supported by the CSIR (Council of Scientific and Industrial Research) -fellowship and also by short-term EMBO Scientific Exchange Grant to carry out part of this work at Swiss TPH. DTP acknowledges the fellowship from the Council of Scientific and Industrial Research for the support. This work was supported by MoE-STARS-2/2023-0249 and CRG/2023/001838 from the Government of India to KK. We are thankful to the NGS facility, and microscopy facility at IISER Pune for their support. SM acknowledges Anusandhan National Research Foundation for the J.C. Bose Fellowship (JCB/2023/000016), Department of Science and Technology for a DST-FIST grant (SR/FST/LS-II/2017/97 to the Department of Biological Sciences, IISER Mohali), and SKR acknowledges the fellowship from the Council of Scientific and Industrial Research. The funders had no role in study design, data analysis, decision to publish, or preparation of the manuscript.

## Authors’ contributions

DVM designed, performed, and analyzed data presented in this study. DP designed, performed experiments and analyzed data related to the FRAP assay in the cells and HiC sequencing. IN designed (plasmid design and cloning) CRISPR/Cas9 strategy for the transfections. IN and DVM contributed to the transgenic line generation (transfections and cell line maintenance) for PfHP1. TSV coordinated and supervised the transgenic lines generation. SKR and DVM designed, performed experiments and analyzed data for the *in vitro* phase separation assays and it was coordinated and supervised by SM. PG designed, performed experiments and analyzed data for the single molecule DNA tethering experiments under the guidance and supervision of MG. DVM and KK wrote the manuscript. KK planned, coordinated, and supervised the project. All authors read and approved the final manuscript.

## Conflict of Interest

The authors declare that they have no conflict of interest.

## Supplementary Figures Legends

**Supplementary Figure 1: *var* introns have a dynamic heterochromatin profile.**

A) Examples of *var* gene with active modifications showing ChIP-seq profile (peak called bedgraph file) of H3K9ac, H4K8ac, H3K79me2, PfHP1, PfSir2A and H3K36me3 along with the consensus motifs identified in Avraham et al. 2012, *var* intronic sequences. B) without active modifications in the intronic motif. Have shown ChIP-seq profile (peak called bedgraph file) of H3K9ac, H4K8ac, H3K79me2, PfHP1, PfSir2A and H3K36me3. C) Global ChIP-seq profile (peak called bedgraph file) of PfH2A, PfH2A.Z and PfHP1 over chromosome 7 (The PfH2A and PfH2A.Z ChIP sequencing data is reanalyzed from Bartfai et al., 2010). All ChIP-seq profile are visualised in IGV. D) ATAC sequencing data (bedgraph file) showing chromatin state on *var* gene in different time points (5 hpi, 10 hpi, 20 hpi, 40 hpi), corresponding strand specific RNA sequencing data (separate bedgraph files for + and – strands) are shown, the boxed region indicates the insulator like motif (The ATAC sequencing and RNA sequencing data reanalyzed from Toenhake et al., 2018). This data indicates that *var* introns in general have an open chromatin state and are transcriptionally active.

**Supplementary Figure 2: PfHP1 phase separation assay.**

A) Percentage identity matrix of HP1 homologs based on the multiple sequence alignment of their protein sequences. B) Multiple sequence alignment of HP1 homologs showing conserved residues in CD and CSD, PfHP1 is highlighted by a red arrow. C) Intrinsically disordered regions in the PfHP1 protein and the domain organization is schematically represented at the bottom. NTE-N-terminal extension, CD-Chromodomain, H-Hinge, CSD-Chromo shadow domain, CTE-C-terminal extension. D) Droplet assay of PfHP1 without GFP tag (60 μΜ), labeled with 1% S33C PfHP1-alexa488 in reaction buffer HEPES 20mM, pH 7.4; KCl 150mM (scale bar, 10 μm). E) Droplet assay with PfHP1 (60 μΜ), labeled with 1% S33C PfHP1-alexa488 in reaction buffer and 5% PEG8000 (scale bar, 10 μm). F) Saturation concentration estimation of PfHP1 (60 μΜ) with and without PEG8000. The concentration of the light phase is C_sat_ and the concentrations (in μΜ) are indicated on the top of each bar graph. G,H) FRAP assay with PfHP1 droplets from D and E respectively (means ± SD; n = 5). I) Salt concentration dependence of PfHP1-eGFP (20 μΜ) phase separated droplets. Concentration of KCl used is indicated on the top of each image (scale bar, 10 μm).

**Supplementary Figure 3: PfHP1 phase separation is a tunable process.**

A) Turbidity assay with PfHP1-eGFP and varying concentrations of *P. falciparum* DNA (800 bp). The experiment was done in triplicates and the bar graph represents means ± SD. B) FRAP assay showing fluorescence recovery of droplets with or without DNA (means ± SD; n = 5). C) Droplet assay with PfHP1 (45 μΜ + 1% Alexa 488 S33C_PfHP1) and 601 Widom sequence (concentrations indicted in the image). D) Saturation concentration estimation of PfHP1-eGFP with or without DNA. The concentration of the light phase is C_sat_ and the concentrations (in μΜ) are indicated on the top of each bar graph. E) Droplet assay with PfHP1 (45 μΜ + 1% Alexa 488 S33C_PfHP1) and YOYO-1 labelled 601 Widom DNA (concentrations indicted in the image). F) Droplet assay with PfHP1-eGFP (15 μM) and PfH2A.Z/PfH2A (4 μM). G) Pairwise sequence alignment of PfHP1 and Swi6. Boxed regions indicate the previously reported functional mutations of Swi6 (AcidX: red, loopX: blue, CageX: green, DimerX: yellow). Scale bar for all images is 10 μm.

**Supplementary Figure 4: AT-richness of *P. falciparum* genes.**

A) Median AT-richness calculated for euchromatic gene (blue), heterochromatic genes (red) and *var* genes (green). The heterochromatic genes are the PfHP1 binding gene sequences (intergenic or telomere regions are not included), includes all CVGs, *var* genes and AP2-G. The fasta sequence obtained from PlasmoDB were used for AT-richness calculation. B-E) The percentage AT-content of *var* genes and promoters B) Pf3D7_1240300, C) Pf3D7_0733000, D) Pf3D7_1041300, E) Pf3D7_0412400. The 2 kbp upstream of *var* coding sequence was used as promoter.

**Supplementary Figure 5: DNA tethering experiments.**

A) EMSA showing shift in parasite DNA (20 ng/μl) upon addition of increasing concentration of PfHP1 (0nM, 125nM, 250nM, 500nM, 1μM, 2μM, 4μM and 10μM). B-Protein bound fraction and F-Free DNA. B) Experimental set up for single molecule DNA tethering experiment. A microfluidics system with a syringe was connected to the slide chamber with the channels for DNA tethering. From on end DNA, protein or buffer can be injected and the syringe pump can modulate the flow rate. A top view of the double tethered DNA is shown, in fluorescence imaging the DNA can be tracked to study the dynamics. C) DNA tethering experiment with the 18kbp control DNA (without *var* intronic sequence) and PfHP1-eGFP (1 μM) (scale bar, 1 μm; the red arrow indicates the time point at which the protein was added during the experiment). At the top, a schematic of double tethered 18 kbp DNA; the DNA tethering was done by neutravidin (green) and biotin (orange) interaction; for determining directionality Cy5 (magenta) probe is attached on one end. D) Co-localization of Cy5 tag with DNA molecule for determining the directionality (19kbp control DNA and PfHP1-eGFP (1 μM) scale bar, 1 μm). E) Percentage of double tethered DNA showing at least one punctum for PfHP1 wildtype and mutants E-A and K-A (different concentrations are used, indicated in the table). F) Proposed model for DNA condensation by PfHP1. The white lines represent DNA and highlighted red regions are extremely AT-rich (∼90%) stretches in the genome like *var* promoters/intron. The nucleosomes with different composition are given at the bottom legend. The heterochromatin formation is a multi-step process where intermolecular interaction with AT-rich DNA and PfHP1 helps in nucleation, and forms co-condensates. The DNA-PfHP1 condensates capture more DNA and starts the compaction. The multivalent interaction of PfHP1 with itself and H3K9me3 helps in the heterochromatin spread and LLPS droplet formation. Further growth and maturation of multiple droplets by fusion leads to heterochromatin domain formation. Additionally, the boundary of these droplet/s might be maintained by PfH2A.Z.

**Supplementary Figure 6: Ectopic expression lines of PfHP1 mutants.**

A) A schematic showing CRISPR-Cas9 strategy for inducible ectopic expression of PfHP1 mutant lines in the *cg6* loci and the primers used for the PCR validation are indicated by purple arrows. The PfHP1 is cloned under cam promoter and is tagged with GFP. The glmS ribozyme would degrade the PfHP1-GFP mRNA in the presence of glucosamine. The Sheild-1 will FKBP-DD to stop the proteasomal degradation of the PfHP1-GFP protein. B,C) PCR validation of ectopic expression lines, the primers used and the expected sizes are given in the table. Three sets of primers were used for the validation. The first set is cg6 UTR primers that should give an amplification in all cell lines including 3D7, but different sizes. The second set of primers were cg6 5’ UTR forward and PfHP1 reverse; this will only give a band if there is a knock in of PfHP1 in the *cg6* locus. The third set of primers were cg6 5’ UTR forward and cg6 inside reverse primers; this should only give bands if the knock-in didn’t happen and wild-type locus is unchanged. The well numbers 1 and 12 have size markers. D) Western blot validation of the lines using anti-PfHP1 antibody (1:5000 dilution) (18). The cell lines were treated with 1.25 μM Sheild-1 for one generation before the imaging. The lanes are 1-ladder, 2-PfHP1-GFP (– Sheild-1), 3-PfHP1-GFP (+ Sheild-1), 4-W29A (– Sheild-1), 5-W29A (+ Sheild-1), 6-L254D (– Sheild-1), 7-L254D (+ Sheild-1), 8-K-A (– Sheild-1), 9-K-A (+ Sheild-1), 10-L254D (– Sheild-1), 11-L254D (+Sheild-1), 12-3D7. The green arrow indicates PfHP1/mutants with GFP tag and the red arrow indicates the untagged endogenous PfHP1. E) Live cell fluorescence imaging of schizonts of the wild-type or mutant lines treated with 1.25 μM Sheild-1. DNA has been stained with Hoechst (scale bar, 5 μm). F) Normalized mean tag density from the PfHP1-GFP/mutant-GFP ChIP-seq, plotted against gene length of *var* genes. The data is normalized to the number of reads. G) Wildtype and PfHP1 mutant binding profile on *var* loci. ChIP seq was done using anti-GFP antibody. The data was visualized in IGV by using input normalized bedgraph files.

**Supplementary Figure 7: Validation of conditional mutant lines.**

A) A schematic showing knock-in of PfHP1 mutant lines in the endogenous *pfhp1* locus, the primers used for the PCR validation are indicated. The red asterisk indicates the stop codon and blue triangle indicate the sera2 intron: loxP. The thick black line is for hrp2 3’ terminator sequence. B) PCR validation of conditional mutants and wild-type of PfHP1. The forward and reverse primers are used as indicated in the schematic; the expected sizes are 1906 bp for the wild-type parent line 3D7/1G5/DiCre, 4879 bp for the transgenics without rapamycin treatment, 2599 bp for the lines with rapamycin treatment which recombines the loxP sites. The loading order is 1-3D7/1G5/DiCre, 2-PfHP1/DiCre (-RAPA), 3-PfHP1/DiCre (+RAPA), 4-PfHP1^W-A^/DiCre (-RAPA), 5-PfHP1^W-A^/DiCre (+RAPA), 6,7-1kbp ladder, 8-PfHP1^K-A^/DiCre (-RAPA), 9-PfHP1^K-A^/DiCre (+RAPA), 10-1kbp ladder, 11-PfHP1^E-A^/DiCre (-RAPA), 12-PfHP1^E-A^/DiCre (+RAPA), 13-PfHP1^L-D^/DiCre (-RAPA), 14-PfHP1^L-D^/DiCre (+RAPA). C) Sanger sequencing validation to indicate the insertion of mutations in the *pfhp1* locus. The PCR product from the amplification in the supplementary Figure 8B was used for Sanger sequencing. The mutated nucleotides and amino acids are highlighted in yellow. D) Sanger sequencing validation of rapamycin induced recombination. The recombined locus will have 5’ UTR followed by a sera2 intron: lox P and re-codonized PfHP1 mutant (in this case L-D).

**Supplementary Figure 8: Live fluorescence imaging of conditional mutant lines.**

A) Live cell fluorescence image showing mScarlet-I signal of PfHP1 wild-type when there is no rapa treatment (scale bar, 5 μm). The PfHP1 mutant lines will also produce a wild-type mSca-I tagged protein when there is no rapamycin. B) Live cell fluorescence imaging of K-A mutant line, 4 days after the rapa treatment (scale bar, 5 μm). C) Bright field images of E-A mutant showing sexual conversion upon rapa treatment, images were taken 8 days after the rapa treatment. The E-A line with DMSO treatment was used as a control.

**Supplementary Figure 9: PfHP1 point mutations lead to hyperactivation of *var* genes.**

A,B) Screenshots of normalized RNA sequencing reads of + and – strands on the *var* genes. The blue color indicates the + reads coming from the *var* gene and the pink color indicates the – reads coming from the *var* genes. The mutants are labelled on left and wild-type (bottom most tracks) is the control in all conditions. The RNA-seq was done in triplicates for representation purpose replicate-1 was used. C-F) Gene ontology analysis of significantly upregulated genes in the PfHP1 mutant line obtained by differential enrichment analysis using DEseq2; the triplicate sequence files of mutant were used and wild-type PfHP1-GFP was used as a control. G) log2FC of *var* genes in the four mutant line from differential enrichment analysis. H) A table summarizing the number of upregulated *var* genes in each mutant base on the differential enrichment analysis log2FC >1 and p-value <0.05.

**Supplementary Figure 10: PfHP1 point mutations lead to activation of heterochromatin genes.**

A-D) heatmap representation of log2FC of heterochromatic genes other than *var* genes after differential enrichment analysis, log2FC >1, p-value <0.05.

**Supplementary Figure 11: Chromosomal contact maps of PfHP1-GFP and its mutant lines.**

The Hi-C contact matrix visualized in Juicebox, indicating all intra and inter-chromosomal interactions. The transgenic lines are indicated on the top and the chromosome numbers are indicated on the sides of the matrix.

**Supplementary Figure 12: Loss of PfHP1 phase separation leads to distinct changes in chromosomal contacts.**

A) HiC contact map of Chromosome 1 for PfHP1-GFP transgenic lines and its mutants. Along with the log_2_(PfHP1^mut^ - PfHP1-GFP) map generated on Juicebox. B) The loss of interaction of telomeric ends are marked by boxes. C) HiC contact map of Chromosome 4 and chromosome 7 for PfHP1-GFP transgenic lines and its mutants. Log_2_(PfHP1^mut^ - PfHP1-GFP) map generated on Juicebox. D) The loss of contact varies across the mutant lines, PfHP1^K-A^ shows the highest difference with the control line (boxed). There is an increase of centromere-to-centromere interaction across mutant lines compared to the control (black arrow).

**Supplementary Figure 13: Contact frequency maps at different resolutions.**

Due to the digestion with two enzymes (Supplementary File 1), the Hi-C data attained has high resolution. In 50kb, 25kb, 10kb and 5kb, the loss of contact in mutant lines is observed compared to the control PfHP1-GFP transgenic line.

**Supplementary Figure 14: Protein purification profile and PfHP1 antibody validation.**

A) Protein purification profile of PfHP1-His, image from the SDS-PAGE stained with Coomassie Brilliant Blue. B) Western blot of *P. falciparum* nuclear lysate (25 μg) probed with pre-immune sera in 1:200 dilution in 5% milk. C) Western blot of *P. falciparum* nuclear protein lysate (25 μg) probed with anti-PfHP1 antibody in 1:200 dilution in 5% milk. D) Protein purification profile of PfHP1-eGFP (image from the SDS-PAGE stained with Coomassie Brilliant Blue); the lanes are, 1- elution 1, 2- elution 2, 3- elution 3, 4- elution 4, 5- elution 5. The red arrow indicates the PfHP1-eGFP band (∼ 61 kDa).

## Supplementary Videos Legends

**Supplementary Video 1:** PfHP1-eGFP (40 μM) in vitro phase separation assay showing liquid like behavior of the droplets. The droplet fusions can be seen (scale bar, 10 μm).

**Supplementary Video 2:** Single molecule DNA tethering assay with PfHP1-eGFP (1 μM) and 19 kbp (with var intron) DNA. The video is captured with the fluorescence from Sytox orange labelled DNA.

**Supplementary Video 3:** Single molecule DNA tethering assay with PfHP1-eGFP (1 μM) and 19 kbp (with var intron) DNA. The video is captured with the fluorescence from eGFP tagged PfHP1. The video 2 and 3 are captured in same time frame from different fluorescence channels.

**Supplementary Video 4:** Single molecule DNA tethering assay with PfHP1-eGFP (1 μM) and 18 kbp (control) DNA. The video is captured with the fluorescence from Sytox orange labelled DNA.

**Supplementary Video 5:** Single molecule DNA tethering assay with PfHP1-eGFP (1 μM) and 18 kbp (control) DNA. The video is captured with the fluorescence from eGFP tagged PfHP1. The video 4 and 5 are captured in same time frame from different fluorescence channels.

**Supplementary Video 6:** Single molecule DNA tethering assay with E-A-eGFP (1 μM) and 19 kbp (with var intron) DNA. The video is captured with the fluorescence from Sytox orange labelled DNA.

**Supplementary Video 7:** Single molecule DNA tethering assay with E-A-eGFP (1 μM) and 19 kbp (with var intron) DNA. The video is captured with the fluorescence from eGFP tagged E-A mutant. The video 6 and 7 are captured in same time frame from different fluorescence channels.

**Supplementary Video 8**: Single molecule DNA tethering assay with K-A-eGFP (1 μM) and 19 kbp (with var intron) DNA. The video is captured with the fluorescence from Sytox orange labelled DNA.

**Supplementary Video 9:** Single molecule DNA tethering assay with K-A-eGFP (1 μM) and 19 kbp (with var intron) DNA. The video is captured with the fluorescence from eGFP tagged K-A mutant. The video 6 and 7 are captured in same time frame from different fluorescence channels.

## Notes

### Competing Interest Statement

The authors have declared no competing interest.

