## Supplementary Information for "The regulation of virulence gene expression is controlled by phase separation of heterochromatin protein 1 (HP1) in *Plasmodium falciparum*"

**Supplementary Figure 1**

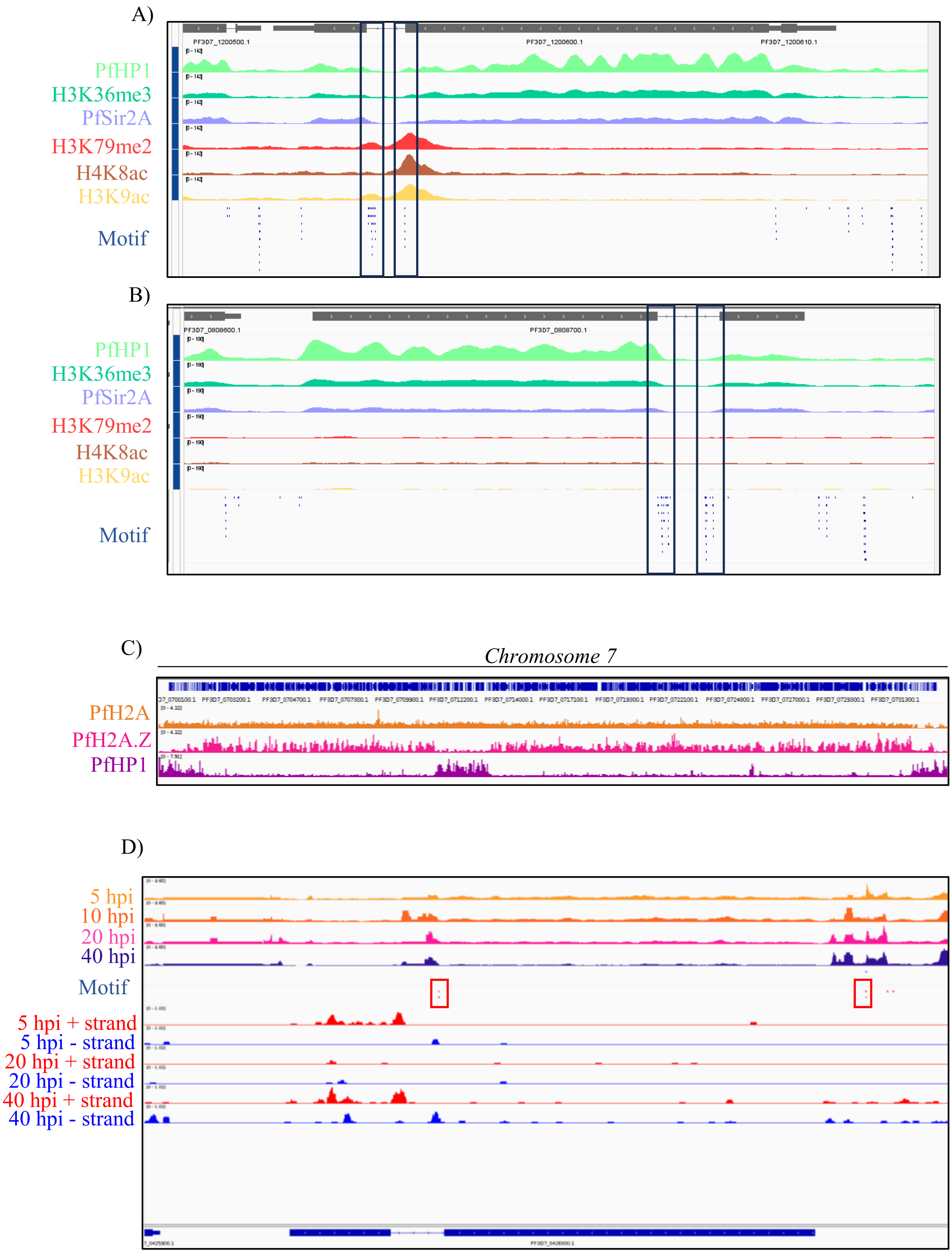

**Supplementary Figure 2**

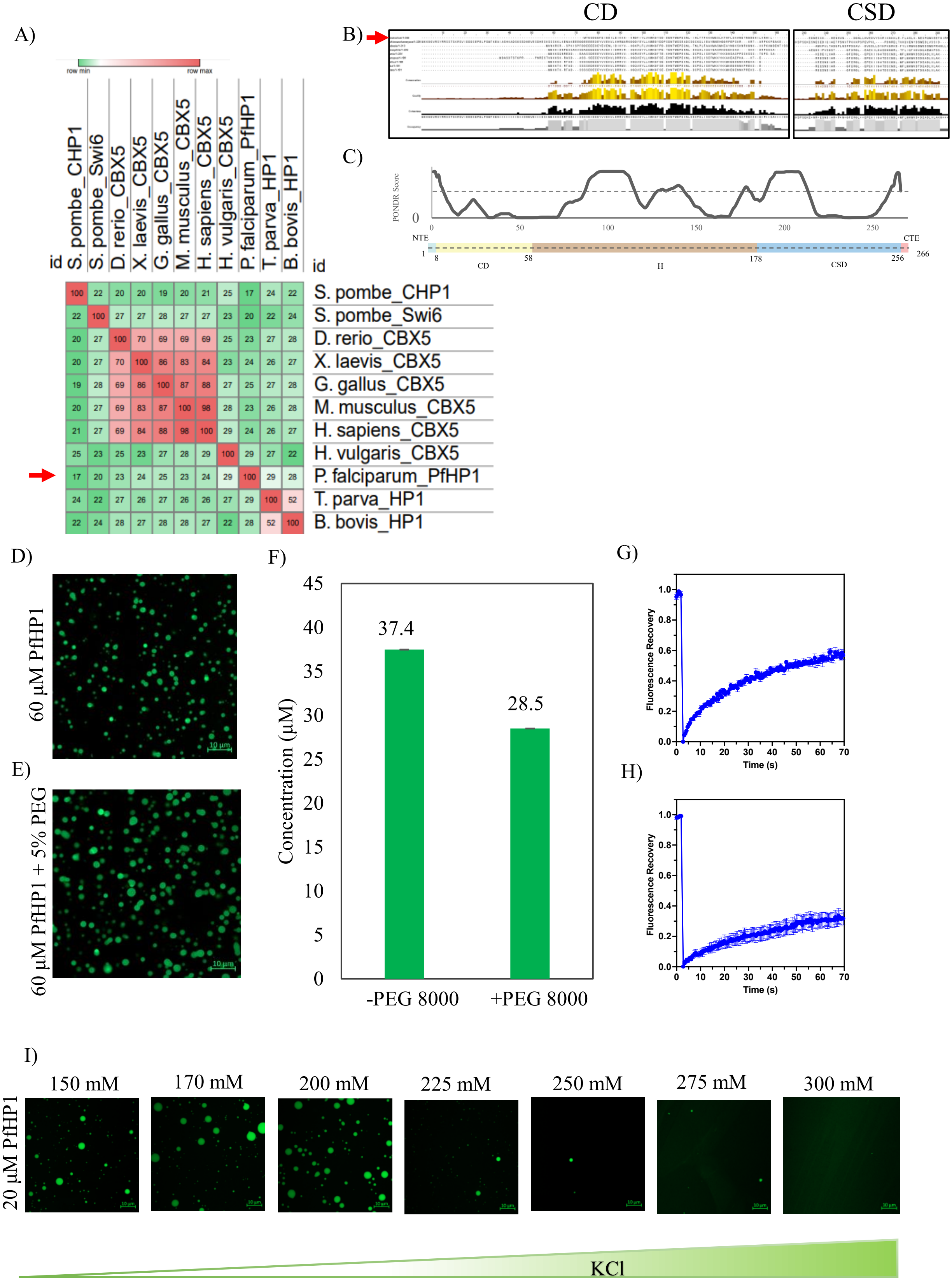

### Supplementary Figure 3

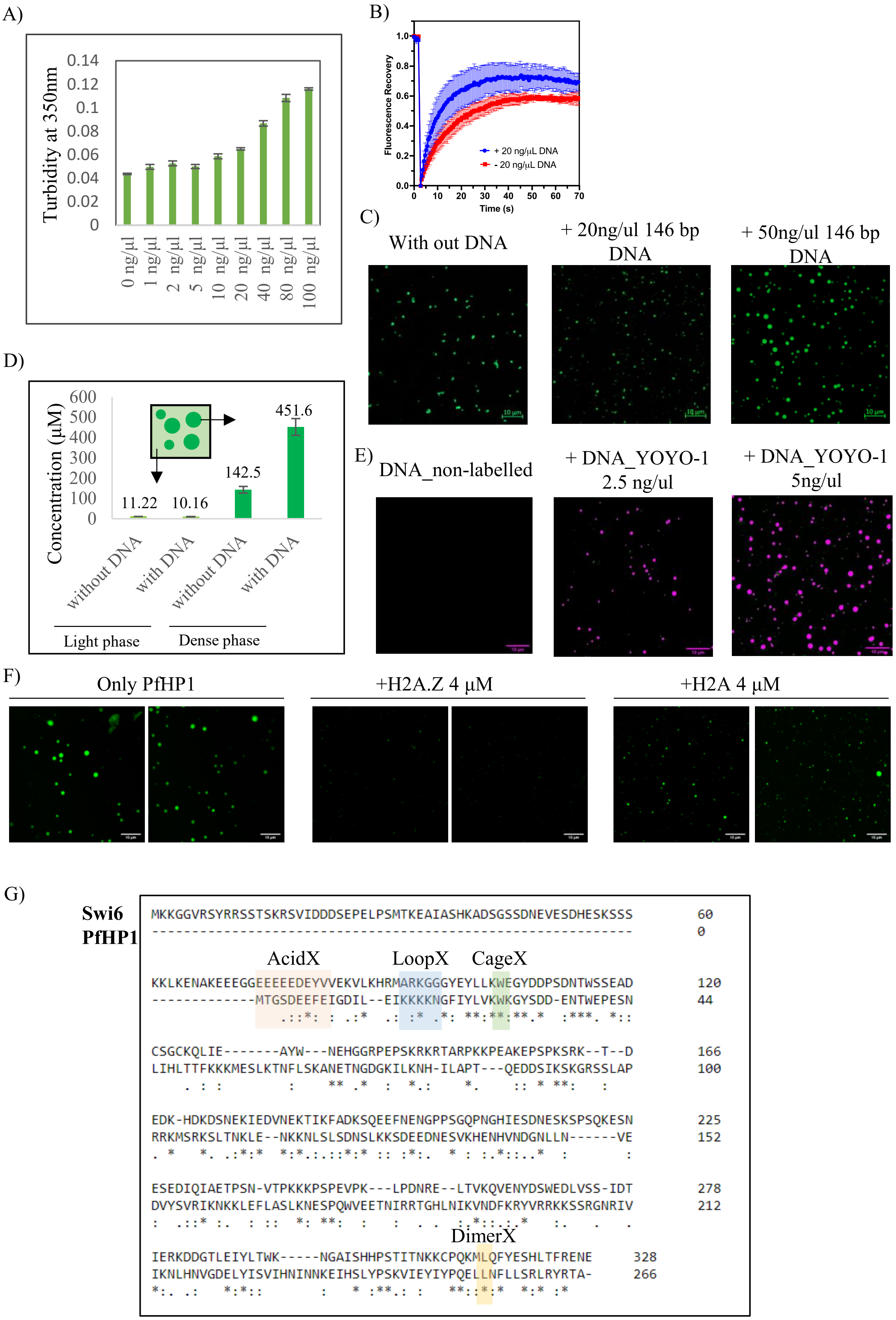

**Supplementary Figure 4**

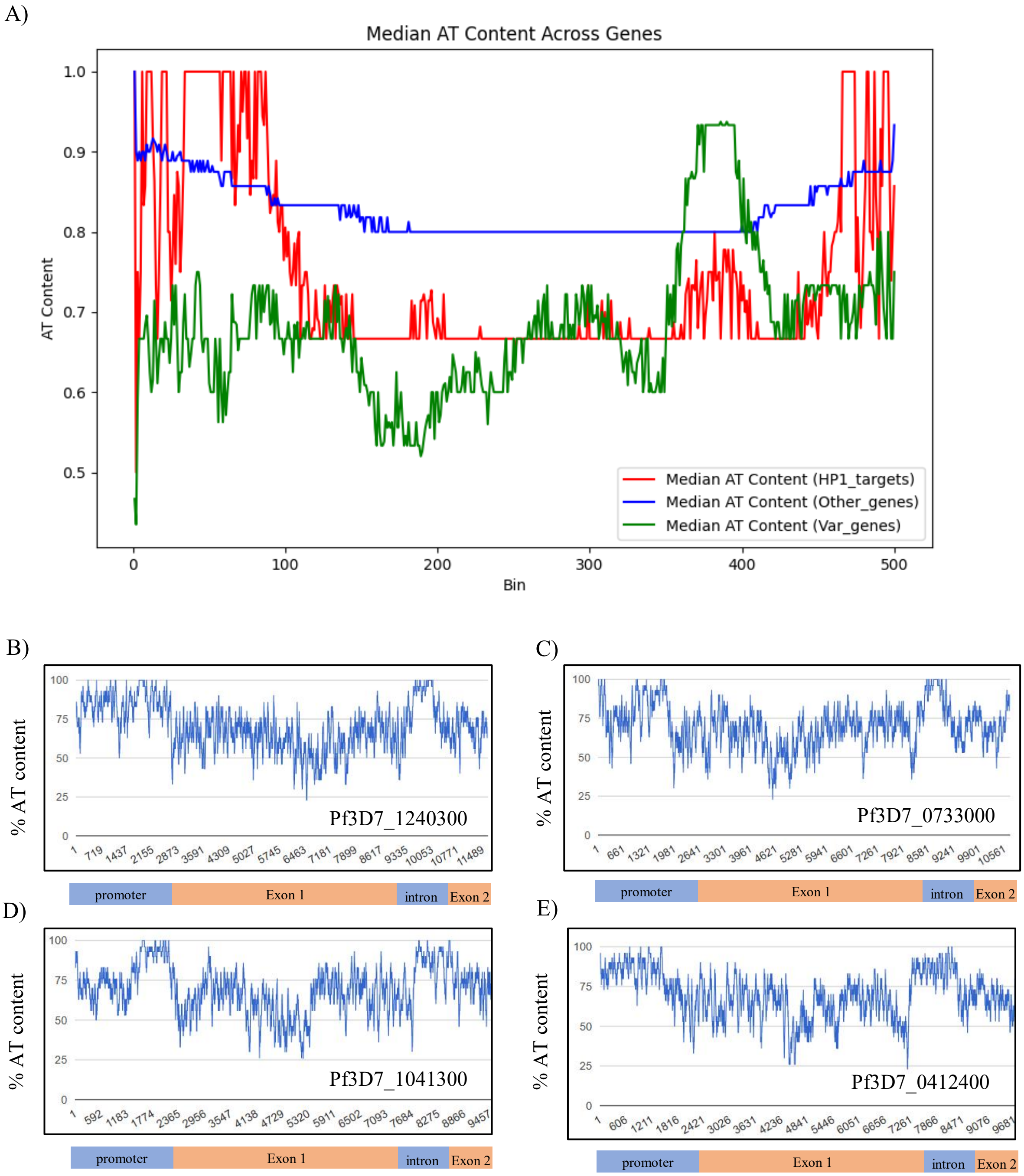

**Supplementary Figure 5**

A)

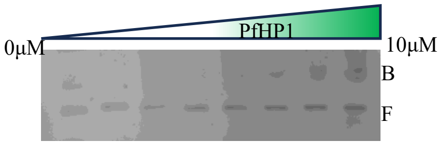


B)

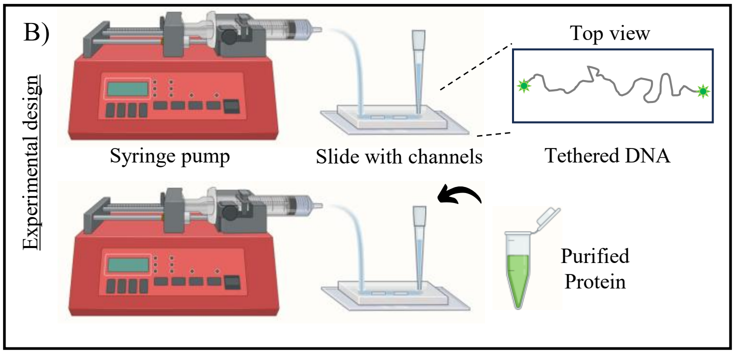


C)

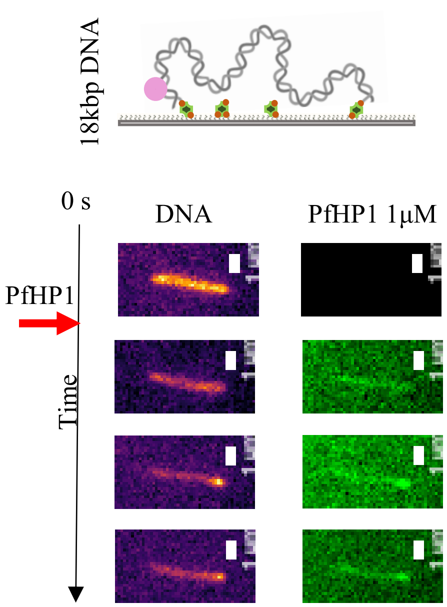


D)

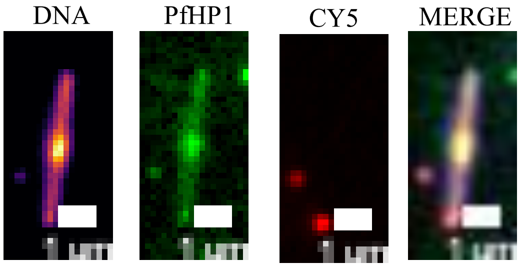


E)

| Condition | % of DNA with puncta |
|---|---|
| WT 500nM | 17.03 |
| WT 1μM | 100 |
| WT 2μM | 100 |
| E-A 500nM | 0 |
| E-A 1μM | 92.85 |
| E-A 2μM | 100 |
| K-A 500nM | 0 |
| K-A 1μM | 0 |
| K-A 1μM | 100 |

F)

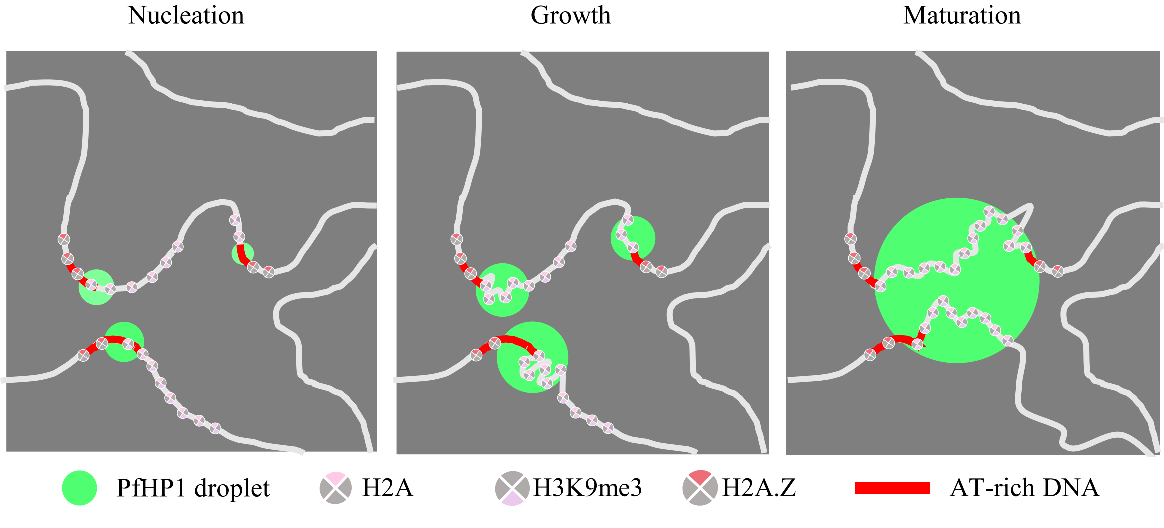

**Supplementary Figure 6**

A)

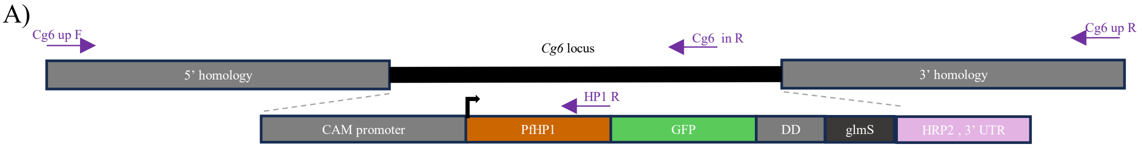


B)

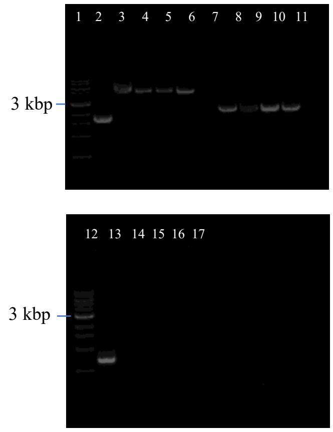


C)

| no. | sample | Primers | Expected size(approx.) |
|---|---|---|---|
| 2 | Control 3D7 | Cg6 up F- cg6 down R | 1.7 kbp |
| 3 | PfHP1-GFP (clone 1) | Cg6 up F- cg6 down R | 5.7 kbp |
| 4 | PfHP1-GFP (clone 2) | Cg6 up F- cg6 down R | 5.7 kbp |
| 5 | W29A-GFP | Cg6 up F- cg6 down R | 5.7 kbp |
| 6 | L254D -GFP | Cg6 up F- cg6 down R | 5.7 kbp |
| 7 | Control 3D7 | Cg6 up F- HP1 R | - |
| 8 | PfHP1-GFP (clone 1) | Cg6 up F- HP1 R | 1.8 kbp |
| 9 | PfHP1-GFP (clone 2) | Cg6 up F- HP1 R | 1.8 kbp |
| 10 | W29A-GFP | Cg6 up F- HP1 R | 1.8 kbp |
| 11 | L254D -GFP | Cg6 up F- HP1 R | 1.8 kbp |
| 13 | Control 3D7 | Cg6 up F- cg6 in R | 950 bp |
| 14 | PfHP1-GFP (clone 1) | Cg6 up F- cg6 in R | - |
| 15 | PfHP1-GFP (clone 2) | Cg6 up F- cg6 in R | - |
| 16 | W29A-GFP | Cg6 up F- cg6 in R | - |
| 17 | L254D -GFP | Cg6 up F- cg6 in R | - |

D)

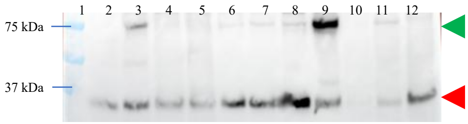


E)

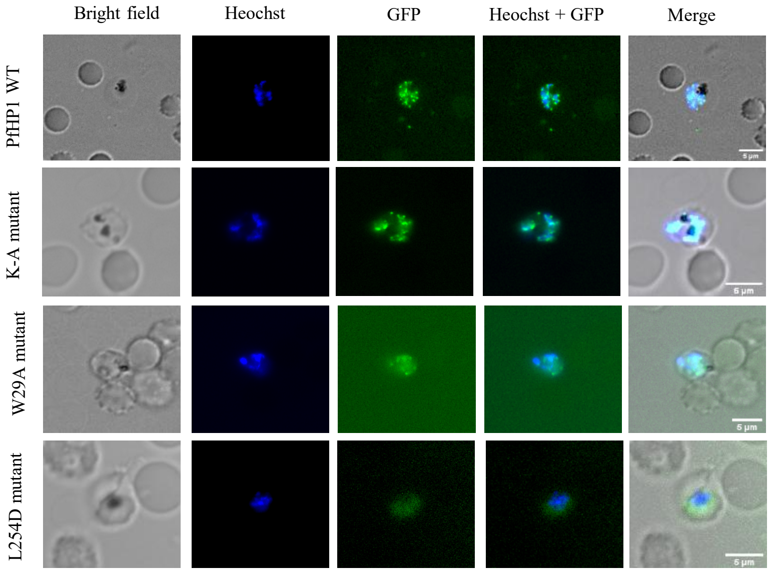


F)

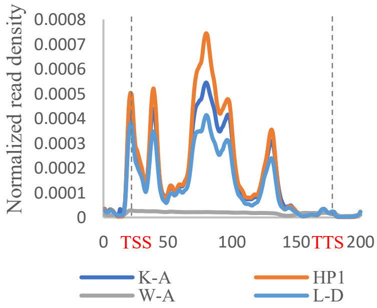


G)

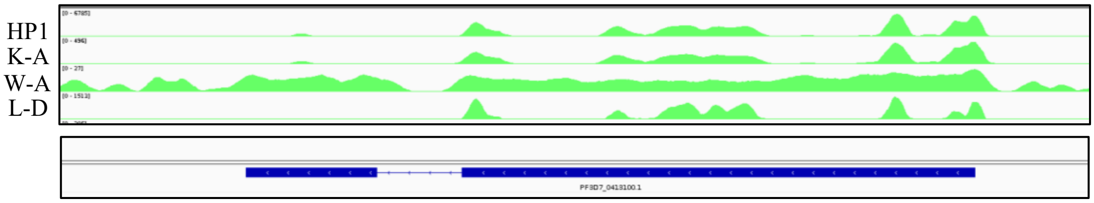

**Supplementary Figure 7**

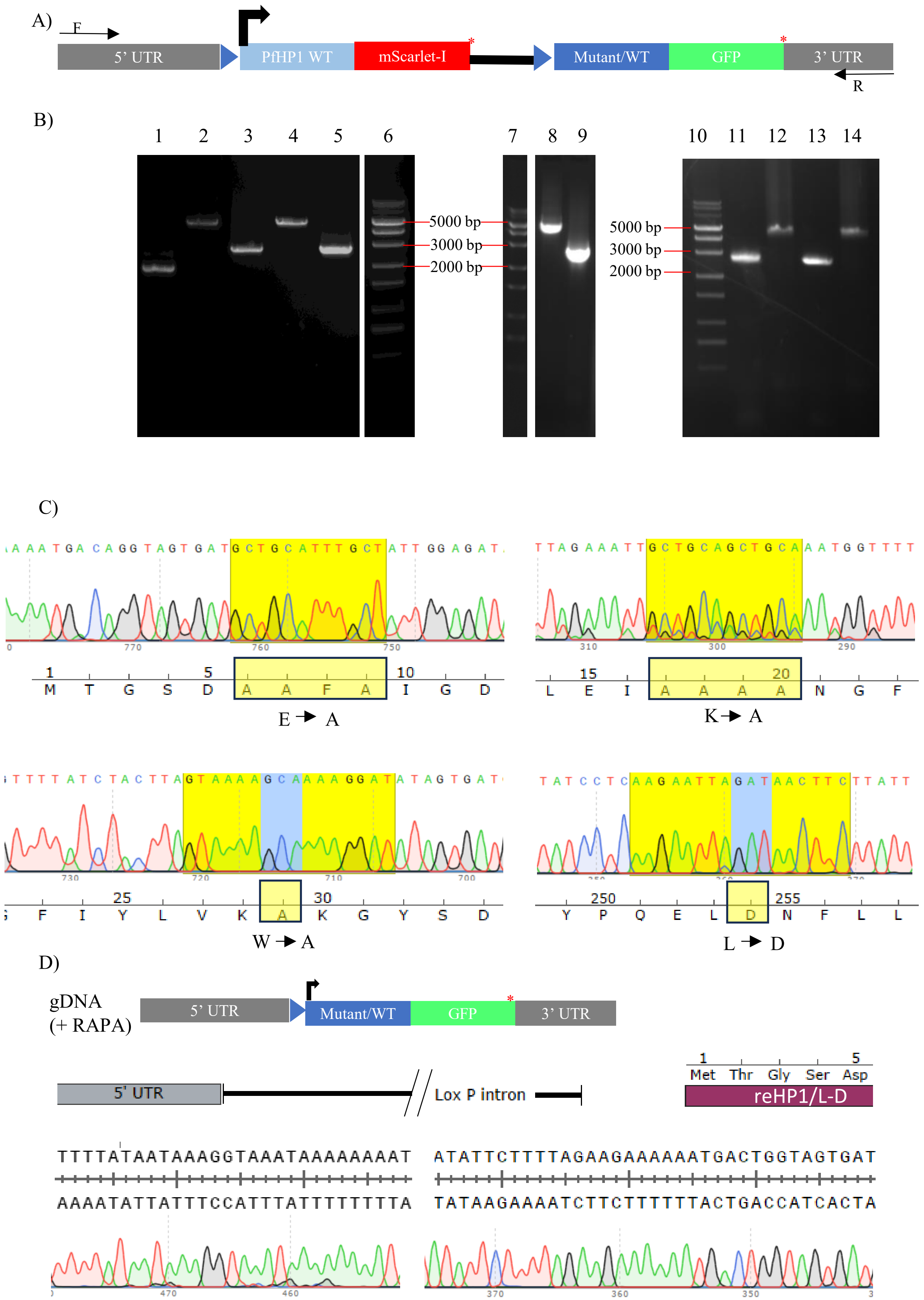

**Supplementary Figure 8**

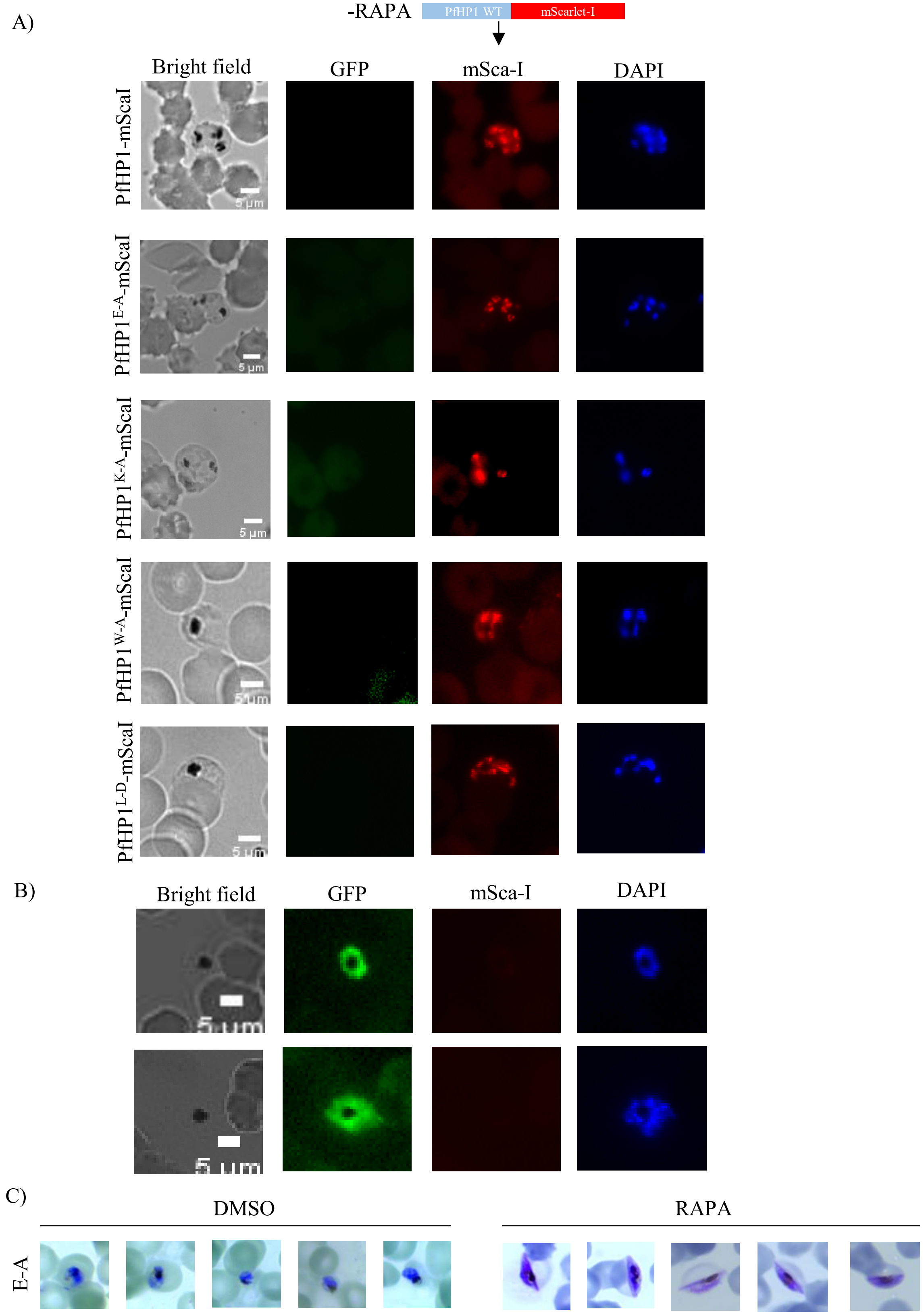

### Supplementary Figure 9

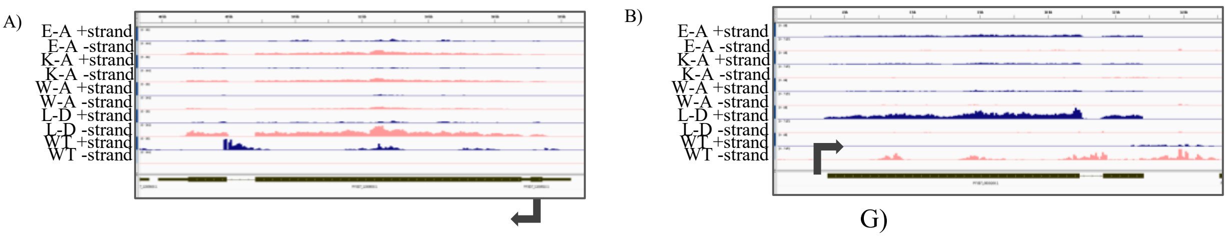


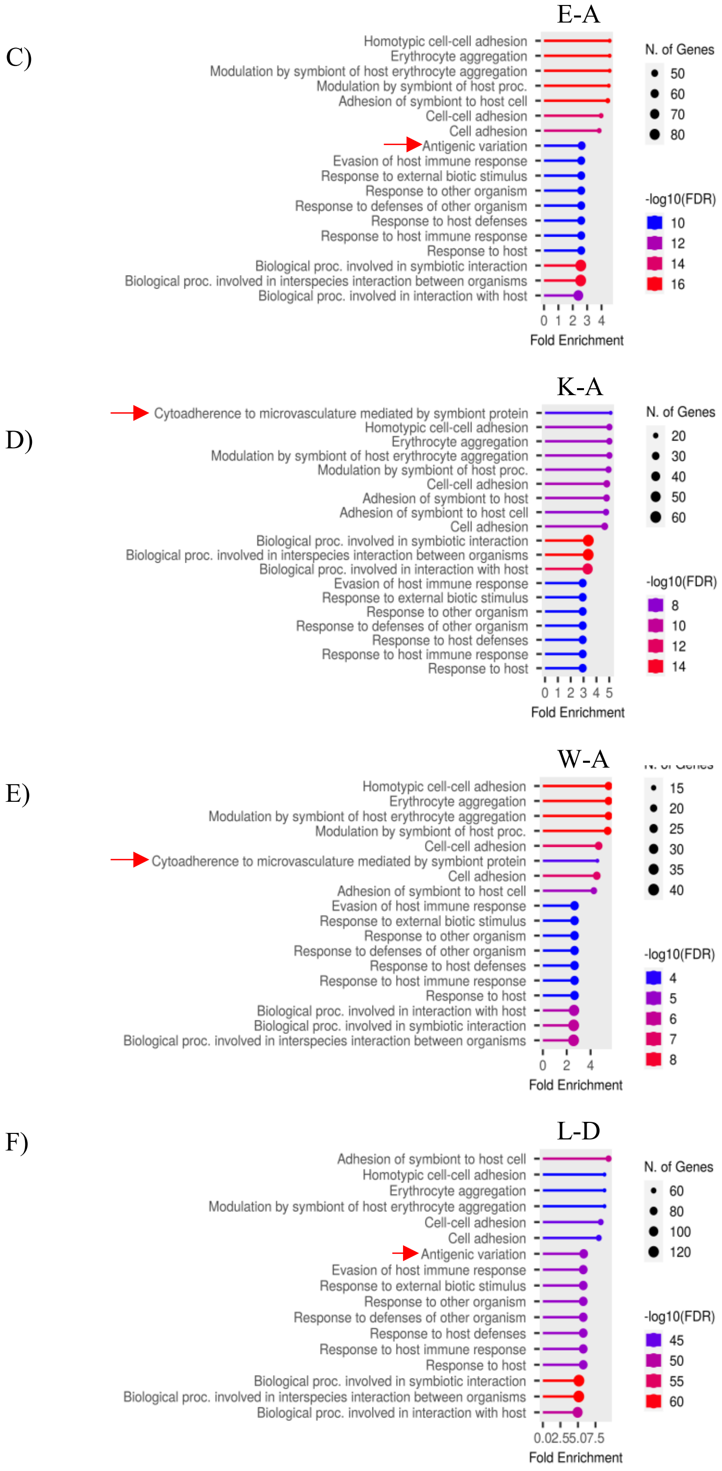


G)

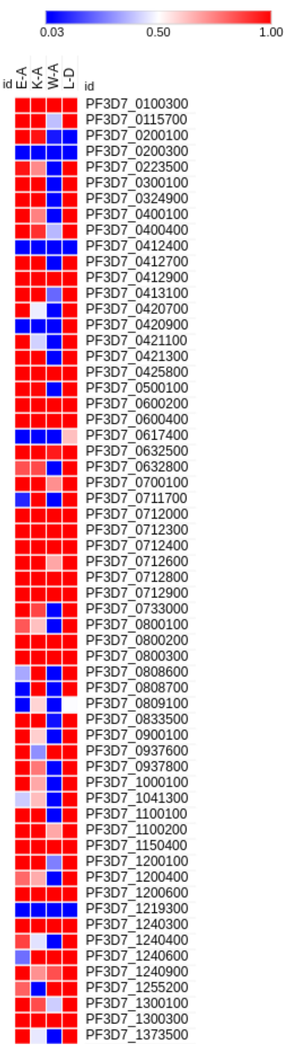


H)

| | E-A | K-A | W-A | L-D |
|---|---|---|---|---|
| # var | 24 | 11 | 10 | 54 |

**Supplementary Figure 10**

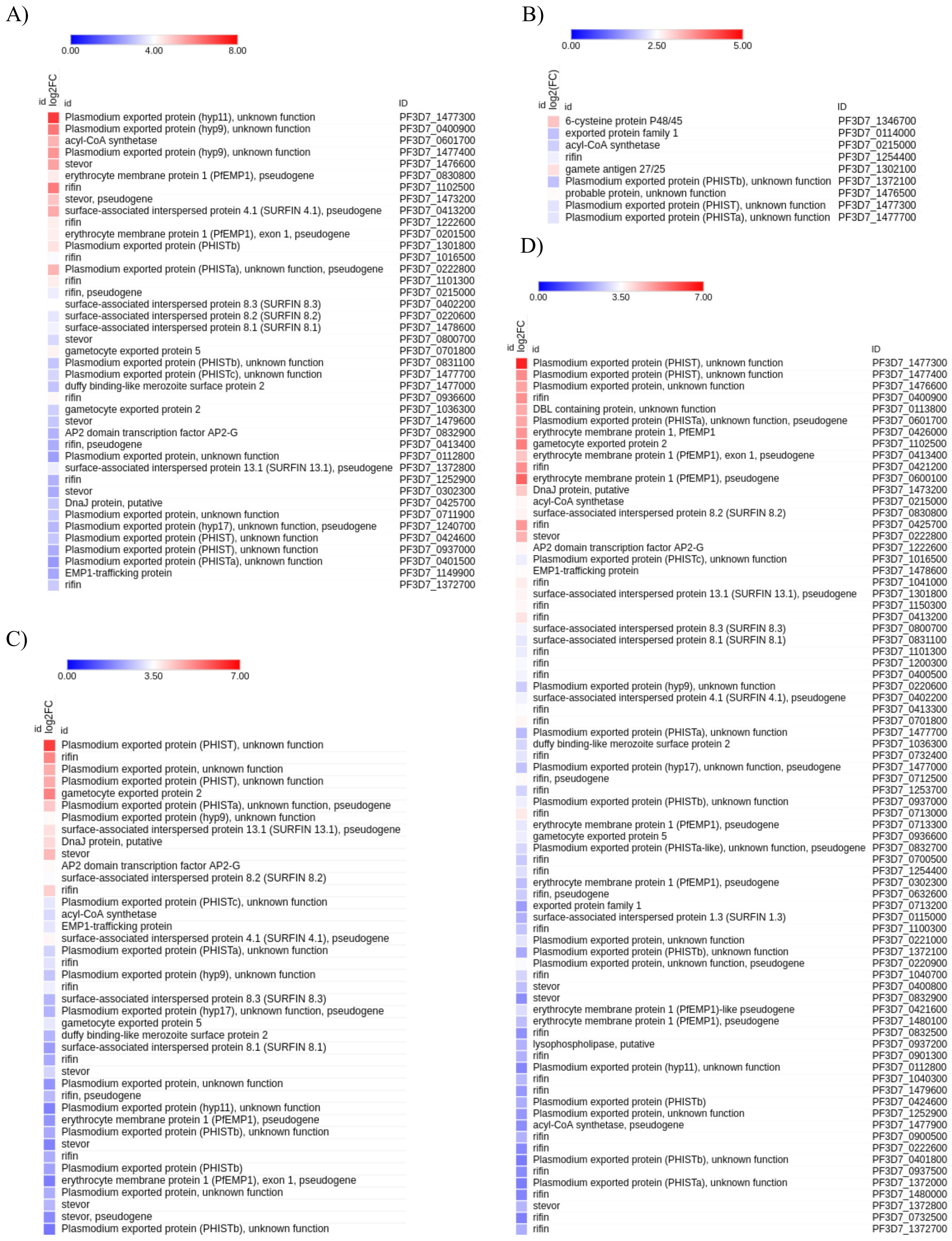

### Supplementary Figure 11

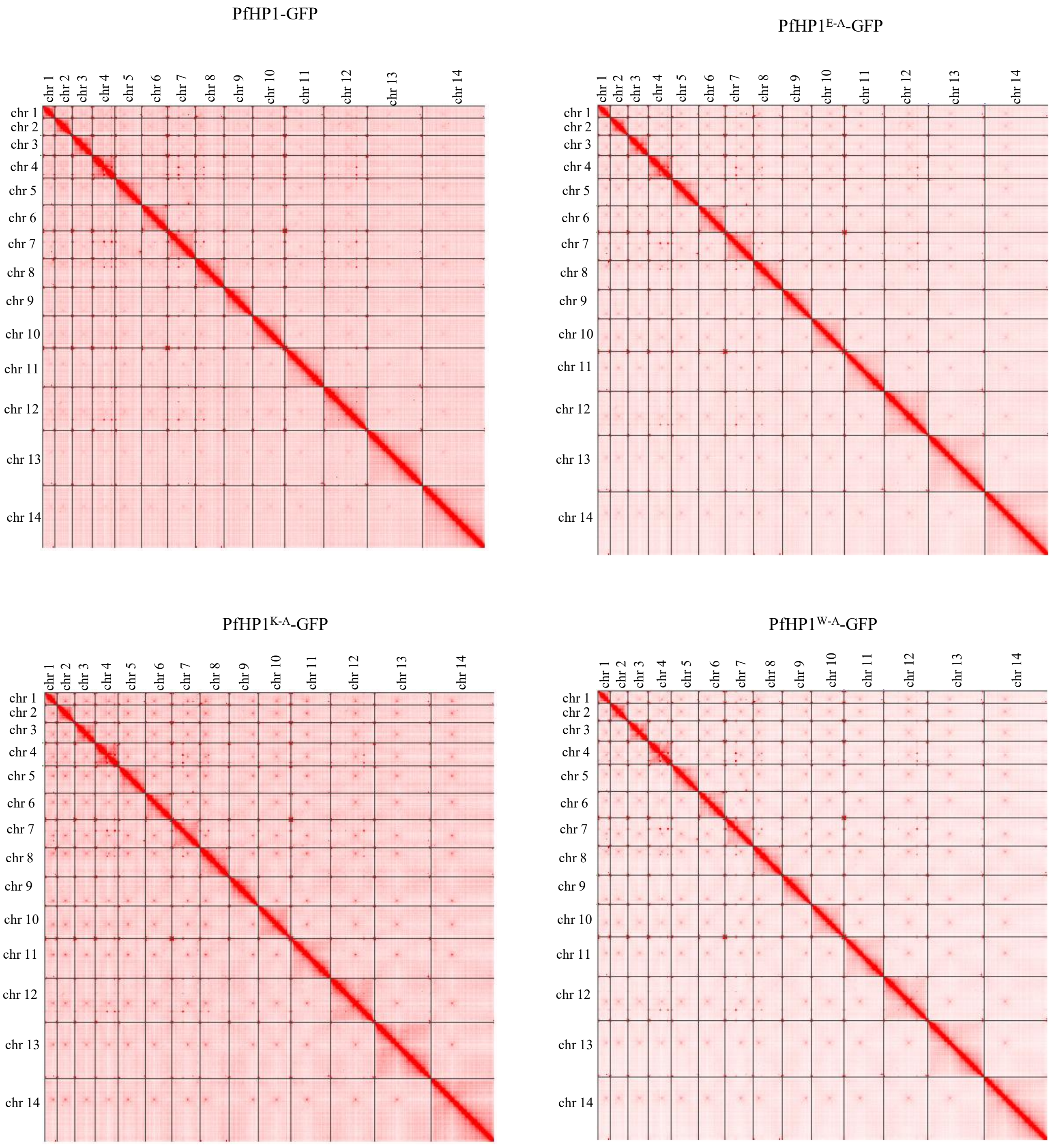

**Supplementary Figure 12**

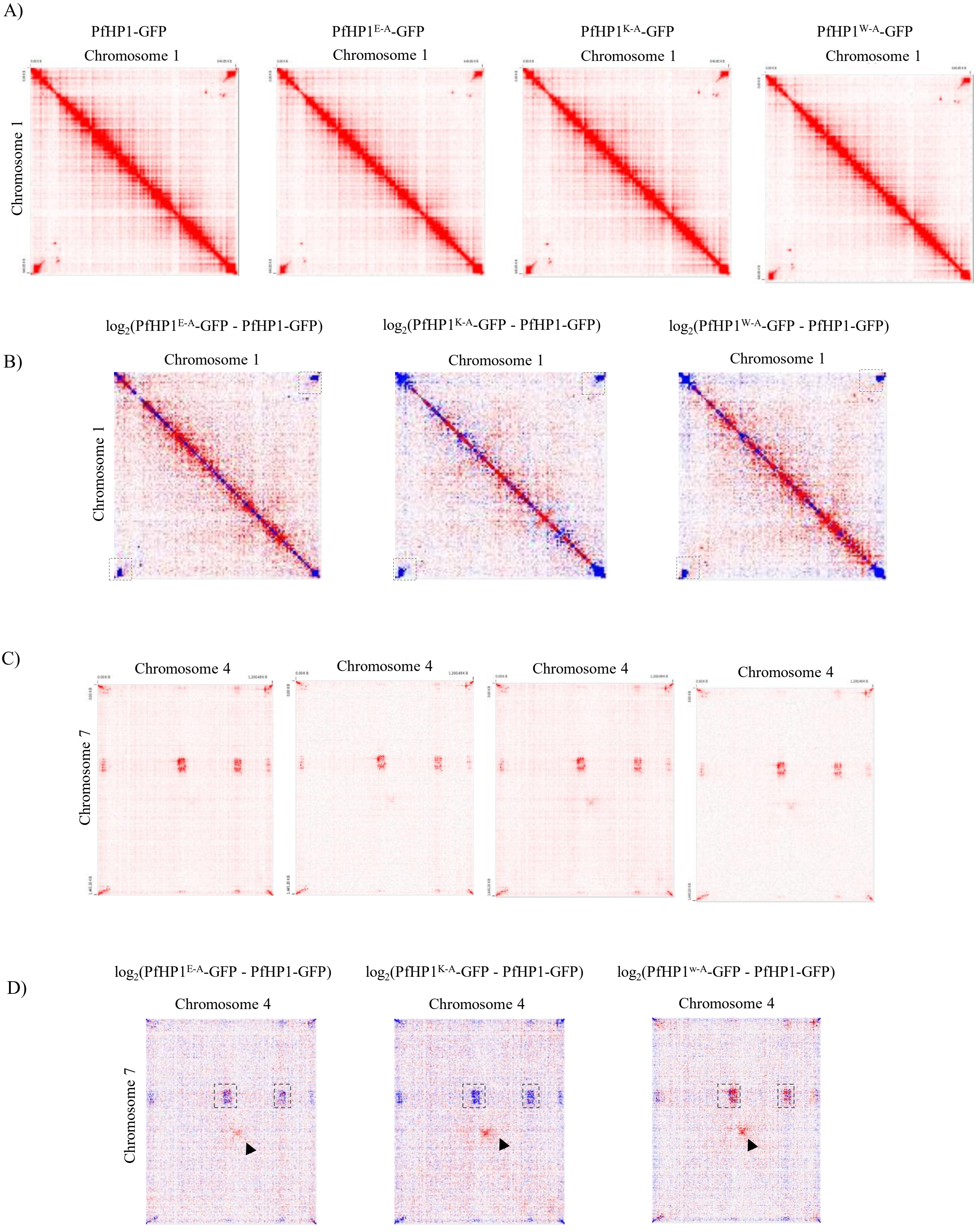

### Supplementary Figure 13

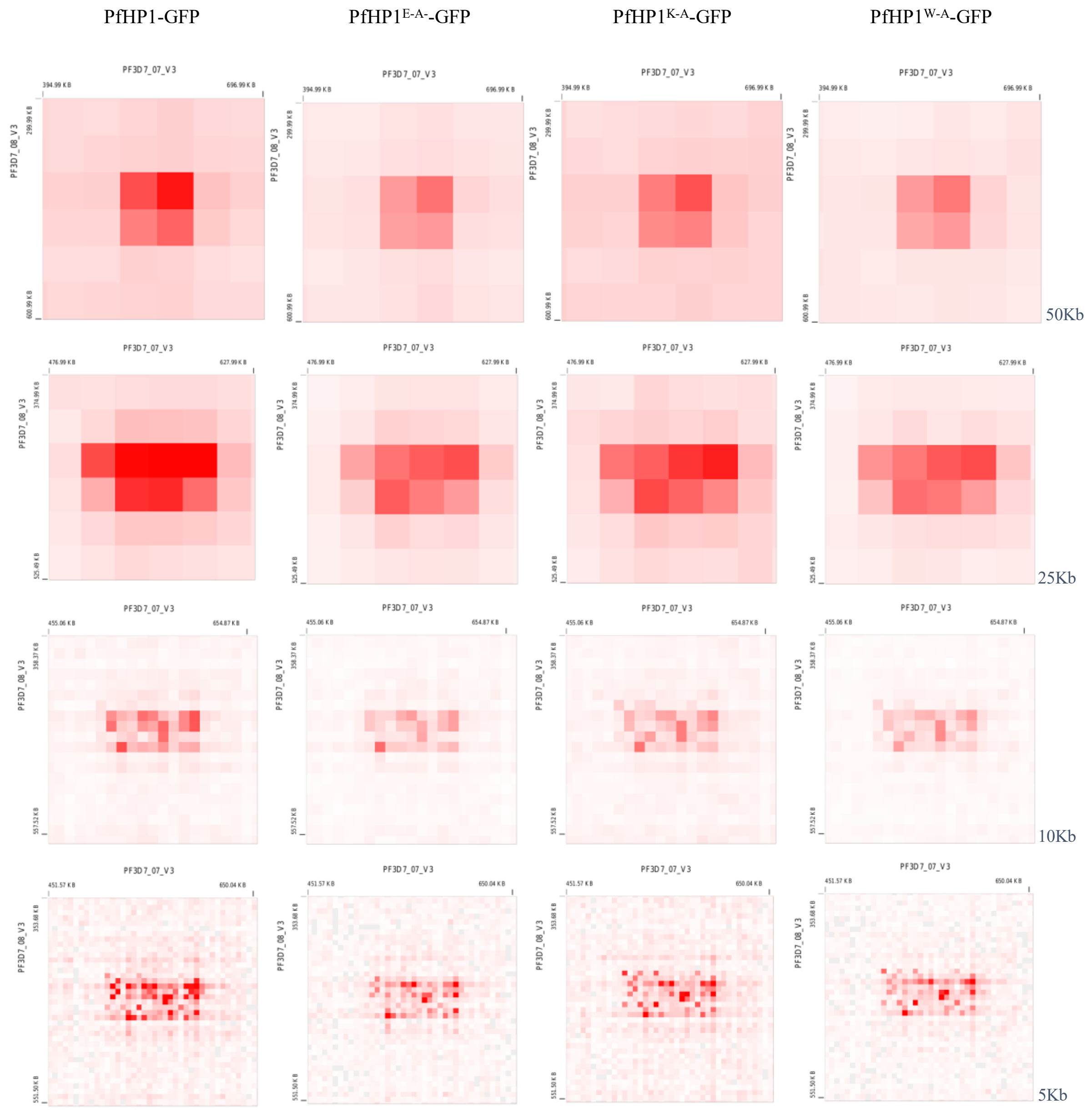

**Supplementary Figure 14**

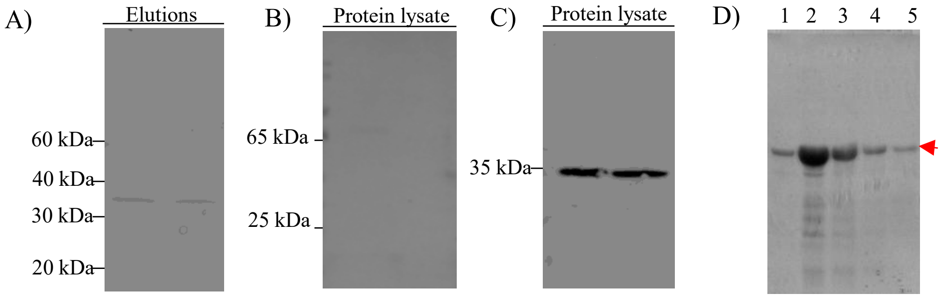
